# Synaptogyrin regulates neuronal activity-induced autophagy to degrade synaptic vesicle components and pathological Tau

**DOI:** 10.1101/2023.07.04.547658

**Authors:** Sergio Hernandez-Diaz, Pilar Martinez-Olondo, Irene Sanchez-Mirasierra, Carla Montecinos, Saurav Ghimire, James D. Sutherland, Rosa Barrio, Sandra-Fausia Soukup

**Affiliations:** Univ. Bordeaux, CNRS, IMN, UMR 5293, F-33000 Bordeaux, France; Université Libre de Bruxelles (ULB), ULB Neuroscience Institute, Neurophy Lab; Bruxelles, Belgium; Univ. Coimbra, Portugal; Center for Cooperative Research in Biosciences (CIC bioGUNE), Basque Research and Technology Alliance (BRTA), 48160 Derio, Spain; Achucarro Basque Center for Neuroscience, Leioa, Spain; Ikerbasque Foundation, Bilbao, Spain

**Keywords:** Autophagy/ *Drosophila*/ Synaptic vesicle degradation/ Frontotemporal Dementia/ Synapse Biology

## Abstract

Synapses are specialized neuronal compartments essential for brain communication. Neuronal communication mostly relies on the adequate supply and renovation of synaptic vesicles that fuse with the plasma membrane and release neurotransmitters in response to action potentials. Autophagy is an evolutionary conserved cellular mechanism essential for homeostasis that can be locally regulated in the neuronal synapse. However, the precise mechanisms controlling synaptic autophagy, especially during neuronal communication and pathological scenarios, remain elusive. Here, we report that neuronal activity and amino-acid deprivation regulate synaptic autophagy via distinct molecular mechanisms. We show that Synaptogyrin, a highly abundant presynaptic protein found in synaptic vesicles, is a novel negative regulator of synaptic autophagy in response to neuronal activity without affecting autophagy induction via amino-acid deprivation. We demonstrate that loss of Synaptogyrin modifies the localization of the autophagy protein Atg9 and boosts autophagosome formation at the synapse. Furthermore, activation of synaptic autophagy by loss of Synaptogyrin, but not by amino-acid deprivation, leads to the degradation of synaptic vesicle components via autophagy. Reducing the levels of Synaptogyrin results in the degradation of synaptic Tau via autophagy and restores autophagy dysfunction observed in a *Drosophila* Tau model of Frontotemporal Dementia (FTD). Our data provide novel and valuable information to understand how autophagy is regulated at the synapse in response to neuronal activity and how this process participates in neuronal (dys)function.

## Introduction

Mature post-mitotic neurons rely on robust quality control mechanisms to maintain cellular homeostasis and communication during their lifetime. Macroautophagy (hereinafter referred to as “autophagy”) is a fundamental homeostatic process that orchestrates the engulfment of diverse cellular components into a growing membrane called autophagosome, that in turn fuses with the lysosome, generating the autolysosome where degradation occurs ^1,2^. Autophagy is crucial in neuronal development ^3^ and during neuronal function and survival ^4^. Much of our knowledge about the regulation of autophagy arises from pioneer work deciphering regulators of autophagy in response to nutrient starvation (e.g. amino acid starvation) ^5–7^. Evidence supports that autophagy is constitutive in neurons ^8,9^ where this process show compartment-specific adaptations ^9–11^. However, little is known about how autophagy is regulated in neuronal compartments in response to stimuli besides starvation such as neuronal activity. Indeed, neuronal activity is reported to activate autophagy, leading to autophagosome biogenesis at the synapse^10,12–16^. Despite their importance, the molecular mechanisms coupling the activation and/or regulation of autophagy in response to these stimuli and to particular cellular processes are still not well understood in neurons and specifically at the synapse ^10^. A shortage of amino acids (herein referred as to amino-acid deprivation) is a classical paradigm to activate autophagy in cells, including neurons, via the activation of cytoplasmic mTORC1, with the aim of maintaining protein homeostasis ^17–19^. Synaptic transmission is an essential component of neuronal communication, a process that relies on multiple proteins coordinating synaptic vesicle biogenesis and recycling, and neurotransmitter release at the presynaptic terminal ^20–22^. The influx of calcium modulates this process in the presynaptic compartment via voltage-sensitive calcium channels ^23–25^. Evidence supports that autophagy takes place and is regulated locally at the presynaptic compartment, where this process may degrade synaptic vesicles ^9,13,26–28^. However, few synaptic proteins are so far reported to participate in the local regulation of autophagy at the synapse ^10,12,29–31^. How neuronal activity and amino-acid deprivation control autophagosome biogenesis at the synapse, and whether these different stimuli orchestrate distinct physiological or pathological processes through specific molecular mechanisms, is vastly unknown.

We report here that amino-acid deprivation and neuronal activity turn on synaptic autophagy using different molecular mechanisms, and these distinct activation pathways could have different functional consequences at the synapse. We show that Synaptogyrin (Syngr), a synaptic vesicle transmembrane protein, controls autophagy at the synapse specifically in response to neuronal activity. In contrast, the previously characterized autophagy protein Endophilin-B (EndoB), solely participates in autophagosome biogenesis in response to amino-acid deprivation and is dispensable for autophagosome biogenesis in response to neuronal activity. Furthermore, we provide evidence that Syngr interacts with Atg9, a core transmembrane autophagy protein that couples the synaptic vesicle cycle and autophagy. We further show that Syngr can regulate the degradation of synaptic vesicle proteins at the presynaptic compartments via neuronal autophagy in contrast to amino acid-induced autophagy. Many neurological diseases are characterized by toxic accumulation of proteins resulting in the degeneration of specific synaptic networks, such as accumulation of Tau in tauopathies ^32–35^, that can also impair the autophagy-lysosomal pathway ^36,37^.

Reduction of Syngr has been proposed as a promising approach to ameliorate synaptic dysfunction caused by pathological Tau accumulation ^32,38^. We show here that expression of pathological mutant Tau results in an increment in the number of autophagic vesicles at the neuromuscular junction synapses of *Drosophila* due to a accumulation of autolysosomes. Reduction of Syngr restores autophagy defects and in turn reduces synaptic levels of the pathological Tau protein at the synapse, providing further mechanistic evidence of how Syngr and synaptic autophagy could ameliorate neuronal defects observed in tauopathies.

## Results

### Synaptogyrin regulates synaptic autophagy in response to neuronal activity

Synapse-enriched proteins such as Endophilin-A, Synaptojanin, AP2, Bassoon, and Endophilin-B (EndoB) participate in the local regulation of autophagy at the presynaptic terminals ^10,12,30,31,39^. However, the number of known regulators of autophagy at the synapses is small, and we are still far from understanding how autophagy is regulated in different compartments of the neuron, especially at the synapse. The presynaptic terminal is a very dynamic compartment that requires constant turnover and recycling of synaptic components, but how autophagy is regulated at the synapse to participate in these processes is still enigmatic. Moreover, we and others showed that neuronal autophagy could be induced by neuronal activity in addition to amino-acid deprivation ^10,12,14,40,41^. Given the close relationship between neuronal activity and synaptic vesicle recycling, we asked whether synaptic vesicle proteins might regulate synaptic autophagy in response to neuronal activity. *Drosophila* Synaptogyrin (Syngr) and its mammalian homologues Synaptogyrin 1 and 3 are highly abundant multispanning transmembrane proteins of synaptic vesicles ^28^ with roles in the regulation of synaptic vesicles size at the neuromuscular junction (NMJ) of *Drosophila* ^42^. At the *Drosophila* NMJ, we found that loss of Syngr (*syngr ^-/-^*) increases the number of autophagosomes at the synapse, detected as Atg8/LC3-positive puncta (Atg8-mCherry clusters) using an unbiased automatic quantification method that we have previously published ^43^ **(Figure 1 A,B and D)**. Conversely, the overexpression of Syngr in motor neurons of *syngr* null mutant animals (*syngr^-/-^;syngr^myc^*) results in a number of autophagosomes comparable to control synapses **(Figure 1 A-D).** Moreover, expressing shRNA against Syngr in motor neurons also leads to increased autophagy levels comparable to autophagy induction via amino-acid deprivation or neuronal activity using the drug Nefiracetam **(Figure S1 A-F)**, indicating that the observed neuronal effects in the *syngr* null mutants are cell-autonomous.

**Figure 1.**
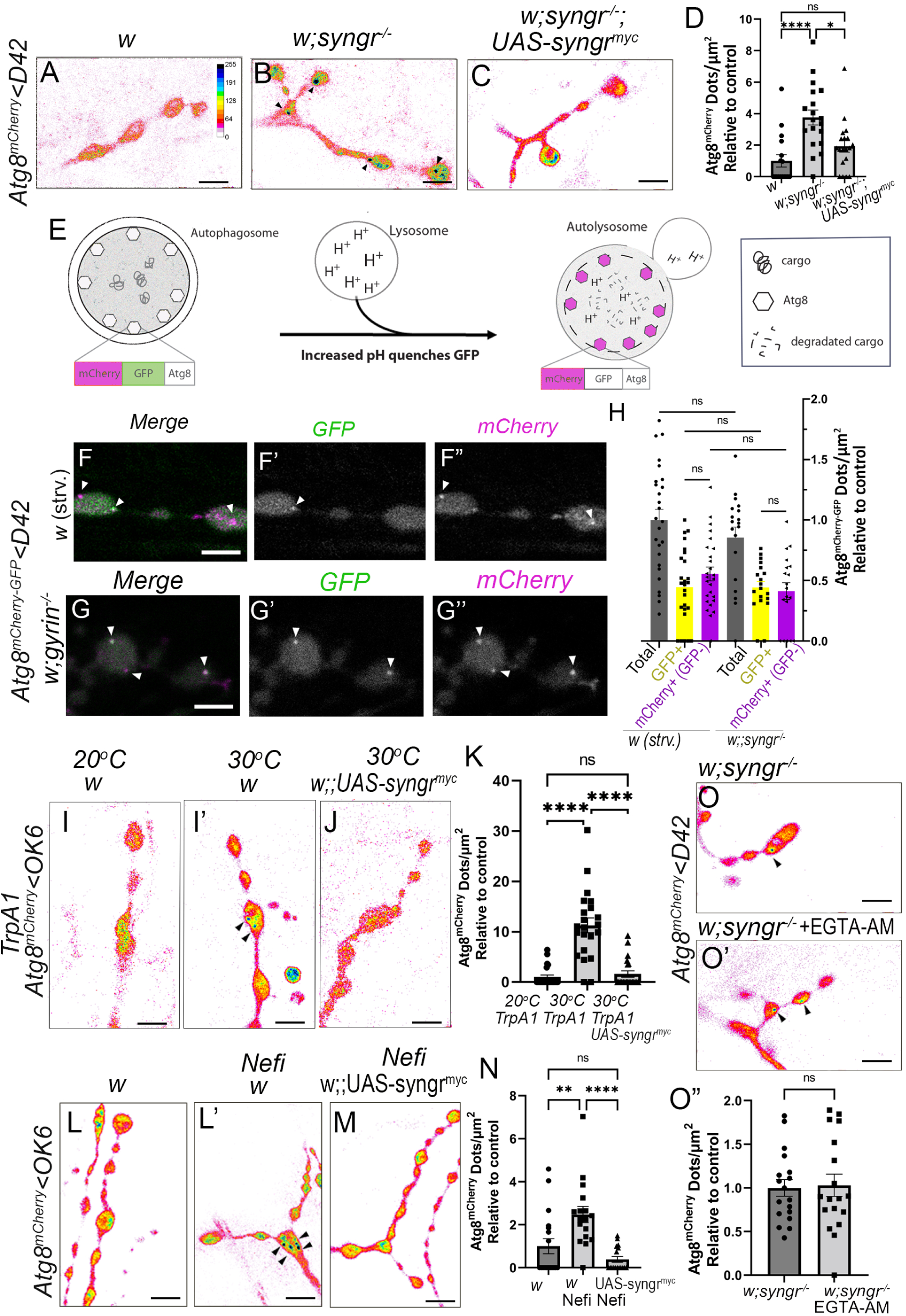
Synaptogyrin regulates neuronal activity-induced autophagy at presynaptic terminals. **(A-C)** Confocal imaging of autophagosomal marker Atg8^mCherry^ at Neuromuscular junction (NMJ) from third instar larvae. Atg8^mCherry^ is expressed using a motor neuron driver (D42-Gal4). Fluorescence intensities are shown using the indicated scale in **(A). (A-B)** NMJ boutons of *syngr* null mutants (*syngr^-/-^*) show increased number of autophagosomes compared to the wildtype (Atg8^mCherry^ positive dots, arrowheads). **(C)** *Expression of Syngr^myc^*in *syngr* null mutants decreases number of autophagosomes comparable to wildtype *syngr* null mutants alone. **(D)** Statistical significances in number of autophagosomes at *syngr* null mutant presynaptic terminals. **(E)** Schematic drawing of autophagosomal assay using the Atg8^mCherry-GFP^ probe. Note that while autophagosomes show both GFP and mCherry fluorescence signals, the GFP fluorescence is quenched in the acidic compartment of the autolysosomes while mCherry fluorescence persists. **(F-G”)** Live imaging of wildtype larvae after induction of autophagy **(F-F”)** and *syngr* null mutants **(G-G”)** expressing Atg8^mCherry-GFP^ to analyze portion of autophagosomes and autolysosomes. **(H)** Quantification of the number of total Atg8 (GFP+mCherry and mCherry alone) dots and the number of white (GFP+mCherry) and magenta (mCherry) Atg8 dots per NMJ area. **(I-K)** Imaging of synaptic boutons labeled with anti-mCherry expressing Atg8^mCherry^ and the TrpA1 thermo sensitive channel (to induce neuronal activity) under control of OK6-Gal4 either at 20°C **(I)** or at 30°C for 30 min **(I’, J)** in wildtype **(I, I’)** or *syngr* null mutant flies **(J)**. Quantification of Atg8^mCherry^ positive dots **(K)** of the indicated genotypes and statistical analysis. **(L-N)** Imaging and quantification of synaptic boutons labeled with anti-mCherry. Imaging of Atg8^mCherry^ dots in wildtype larva with control solution (0,0025% DMSO) **(L)** or Nefiracetam (Nefi) solution, a compound that opens L/N-type Ca2^+^ channels **(L’)** and larvae expressing *Syngr^myc^* in Nefiracetam solution **(M)**. Quantification of Atg8^mCherry^ dots per area **(N). (O-O’)** NMJ boutons of *syngr* null mutants (*syngr^-/-^*) expressing autophagy marker Atg8^mCherry^ incubated with control solution (0,0025% DMSO) **(O)** or the membrane permeable calcium chelator EGTA-AM **(O’)** and quantification of Atg8^mCherry^ dots per area **(O”)**. Scale bars= 5µm. n >19 individual synapses from a minimum of 5 larvae.

To understand if the increased amount autophagosomes in *syngr* null mutant is caused by alterations in autophagosomal biogenesis we analysed the size of autophagosome and did not observe statistical significant size difference of Atg8-positive dots formed after amino acid deprivation, neuronal activity, *syngr* null mutants and *syngr* knock down synapses.**(Figure S1 G)**, Next, we tested whether the increase in the Atg8-positive puncta in *syngr* null synapses arises from increased autophagosome biogenesis or a blockage in autophagosomal flux (failure of autophagosomes to fuse with lysosomes for degradation). For this purpose, we used an autophagic flux marker, consisting of Atg8 tandem-tagged with GFP and mCherry (Atg8^mCherry-GFP^). In this assay, Atg8-positive autophagosomes display both red and green fluorescence, but after fusion with the lysosome, they exhibit only red fluorescence due to GFP quenching in acidic environments **(Figure 1 E)** ^12^. We observe no statistical difference between the quantities of mCherry-GFP and mCherry only puncta in 4h starved wildtype and *syngr* null mutant synapses **(Figure 1 F-H and Figure S1H,I).** Thus, our results show no alteration of autophagy flux in *syngr* null mutant synapses, suggesting that lowering the levels of Syngr increases autophagosome biogenesis. As mentioned above, neuronal activity can promote autophagosome biogenesis at the presynaptic compartment ^10,12,14,15^. Considering that neuronal activity mobilizes synaptic vesicles and that Syngr is a synaptic vesicle protein, we reasoned that Syngr could regulate neuronal activity-induced autophagy. To further test this hypothesis, we took advantage of the genetically expressed thermosensitive ion channel TrpA1 to induce neuronal activation^44,45^ that in turn promotes autophagy induction when animals are shifted to 30^°^C ^10^.

We observed increased amounts of Atg8-positive dots in flies expressing TrpA1 in motor neurons that were shifted to 30^°^C but not in transgenic animals overexpressing Syngr (*syngr^myc^*) **(Figure 1 I-K)**. Using the cation channel agonist Nefiracetam in combination with calcium activates synaptic autophagy by opening L-type calcium channels and therefore mimicking neuronal stimulation ^14,46–48^. While Nefiracetam leads to increased amounts of autophagosomes in wildtype synapses **(Figure 1 L-N** and **Figure S1 D,D’,F)**, in Syngr overexpressing synapses upon Nefiracetam treatment the amount of autophagosomes were significantly lower and comparable to wildtype synapses without Nefiracetam treatment **(Figure 1 M,N),** similarly to the results observed upon autophagy induction using TrpA1, further supporting a role for Syngr as a negative regulator of neuronal activity-dependent autophagy. Reducing the levels of Syngr in addition to Nefiracetam treatment does not increases the number of autophagosomes compared with Nefiracetam or Syngr knockdown individual treatments, likely indicating the existence of a plateau in the induction of synaptic autophagy **(Figure S1 D’,E’,F).** Furthermore, adding a cell-permeable calcium chelator (EGTA-AM) does not block autophagy in *syngr* mutants **(Figure 1 O-O”)**, suggesting that synaptic autophagy in *syngr* mutants is constitutively active and is insensitive to the levels of cytosolic calcium. Taken together, our results support that Syngr is a novel negative regulator of autophagy induction at the presynaptic terminals.

### Synaptogyrin and Endophilin B regulate distinct autophagy pathways at the synapse

Amino-acid starvation leads to increased levels of autophagy in neurons including the *Drosophila* NMJ synapse ^10,12,49^ **(Figure 2 A,A’,C** and **Figure S1 A,A’,C)**. We wondered if Syngr can also regulate autophagy induction via amino-acid deprivation or if Syngr functions specifically in neuronal activity-induced autophagy. Our results show that upon amino-acid starvation autophagy levels are similar in synapses of wildtype and animals overexpressing Syngr (*syngr^myc^)* **(Figure 2 A’,B,C)** and reducing the levels of Syngr in addition to amino-acid deprivation does not affect the amounts of autophagosomes compared with starved control animals **(Figure S1 A,C)**. Rapamycin is a drug that inhibits the nutrient-sensing TOR pathway and in turn activates autophagy ^50^. After 4 hours of rapamycin treatment, autophagosomes accumulate at the wild-type *Drosophila* NMJ **(Figure 2 D,D’,F)**. Accordingly with our previous data, the overexpression of Syngr has no effect on autophagy induction in animals treated with rapamycin **(Figure 2 E,F)**, further supporting that Syngr role is restricted to neuronal activity-induced autophagy.

**Figure 2.**
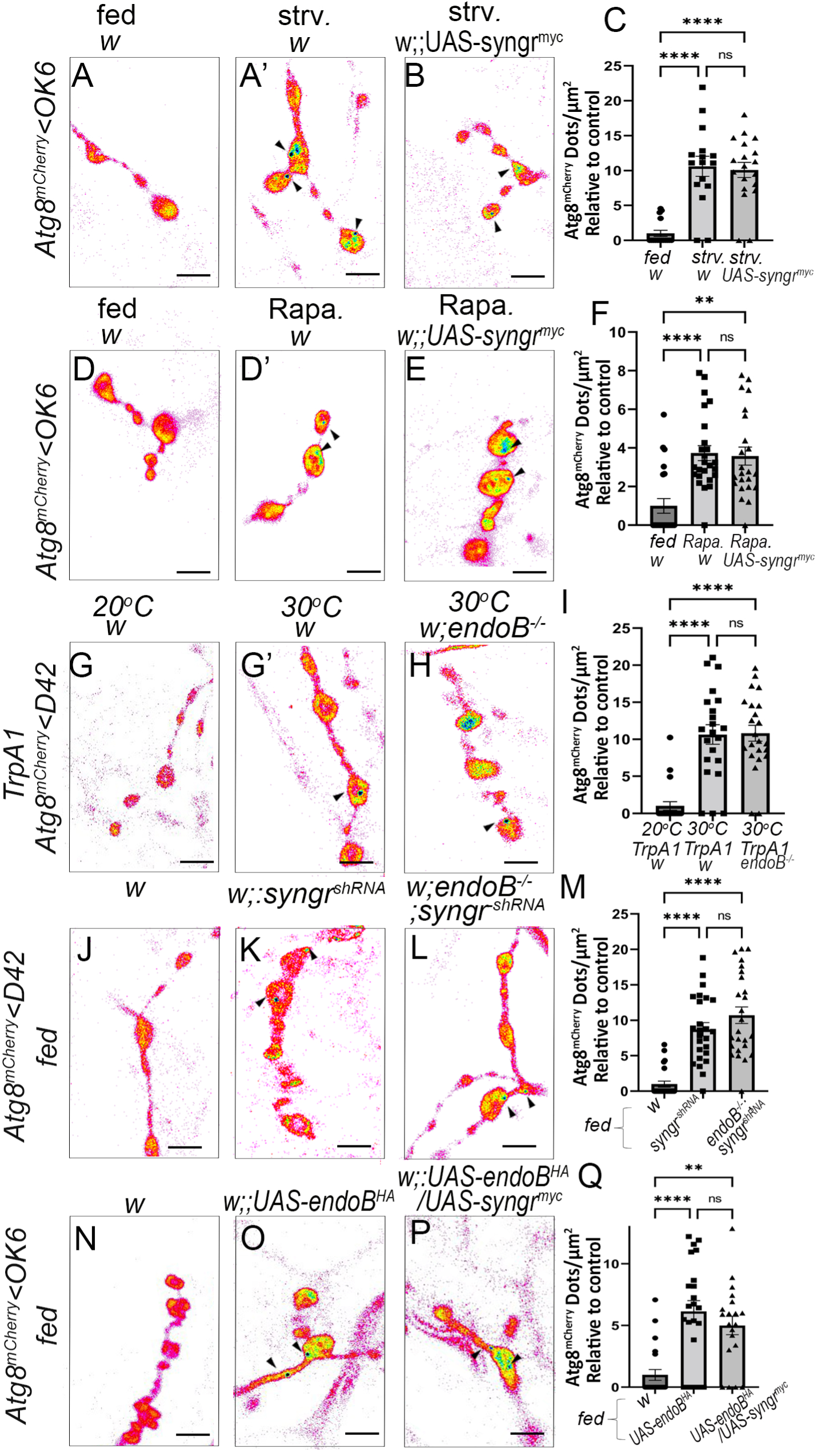
Synaptogyrin and Endophilin-B are required for two molecular different autophagy pathways at the presynaptic terminal. Imaging of fed **(A)** and 4h-starved third instar larvae expressing Atg8^mCherry^ alone **(A’)** or in combination with *syngr* overexpression (*syngr^myc^)* (B) using a motor neuron driver (OK6-Gal4). **(C)** Quantification of Atg8 positive dots show no significant differences between 4h starved control and *syngr* overexpressing larvae (*syngr^myc^).* Imaging of control **(D)** and rapamycin fed third instar larvae expressing Atg8^mCherry^ alone **(D’)** or in combination with overexpression of *syngr* (*syngr^myc^)* (E) using a motor neuron driver (OK6-Gal4). **(F)** Quantification of Atg8 positive dots shows no significant differences between control and Syngr overexpressing (*syngr^myc^)* after rapamycin feeding. Imaging of NMJ boutons expressing TrpA1 channel in wildtype **(G, G’)** and *endophilin-B* null mutant flies (w;*endoB^-/-^)* (H) at 20°C **(G)** or 30°C for 30 min **(G’, H)**. Note that Atg8^mCherry^ dots formation is not significantly different between control and *endoB^-/-^* mutant flies at 30°C **(I)**. Imaging of NMJ boutons of fed wildtype **(J)**, fed flies expressing shRNA against *synaptogyrin* (*syngr^shRNA^)* alone **(K)** or in *endoB* mutant background (w;*endoB^-/-^;syngr^shRNA^)* (L). Statistical analysis of Atg8^mCherry^ dots per area shows that the loss of EndoB has no significant effect in the increase of autophagosomes observed upon knock down of *syngr* (M). Imaging of NMJ synapses from fed third instar wildtype **(N)**, overexpressing *endoB* alone (*endoB^HA^*) **(O)** or in combination with the overexpression of *syngr* (*endoB^HA^*;*syngr ^myc^)* (P) using a motor neuron driver (OK6-Gal4). Quantification of Atg8 positive dots show that the increase in the number of Atg8 puncta observed as result of the overexpression of EndoB cannot be blocked by overexpression of Syngr (Q). Arrows indicate Atg8^mCherry^ dots. Scale bars= 5 µm. (n >19 individual synapses from a minimum of 5 larvae)

The protein Endophilin-B (EndoB) is necessary for autophagy induction in the presynaptic terminal in response to amino-acid starvation ^39^ and **(Figure S2 A-C)** but whether this protein participates in synaptic autophagy in response to neuronal activity is unknown. Interestingly, upon induction of neuronal activity autophagy levels in *endoB* null mutant (*endoB^-/-^*) synapses not significant different to wildtype **(Figure 2 G-I)**. We performed a genetic interaction study to further understand the relationship of Syngr and EndoB in autophagy. Similar to *syngr* mutants, knockdown of Syngr under basal conditions leads to increased amounts of autophagosomes at the synapse **(Figure 2 J,K and M).** Furthermore, knockdown of Syngr in *endoB* null mutant synapses (*endoB^-/-^;syngr^shRN^*^A^) also increases autophagy levels **(Figure 2 L,M).** Moreover, overexpression of EndoB (*endoB^HA^*) activates autophagy at the NMJ of fed animals **(Figure 2 N,O and Q)** and overexpression of syngr does not block autophagy in this condition (*endoB^HA^ ;syngr^myc^*) **(Figure 2 P,Q),** suggesting that EndoB does not function in the same pathway as Syngr. These results indicate that EndoB regulates autophagy in response to amino-acid starvation but not to neuronal activity, while Syngr participates in autophagy in response to neuronal activity but not to amino-acid starvation. Syngr knockdown flies overexpressing EndoB have a similar amount of Atg8mCherry positive puncta compared to Syngr knock down flies alone **(Figure S2 D-G)**. This is not very surprising since Syngr knock down flies with and without Nefiracetam have comparable autophagy levels **(Figure S1 D-F)**. These data further indicate a possible presence of a plateau in the induction of synaptic autophagy that has been reached **(Figure S1 D’,E’,F).** Considering that overexpression of Syngr did not suppress autophagy induction after amino-acid starvation and that mutant *endoB* synapses can normally form autophagosomes upon neuronal activity, our results suggest the existence of at least two molecularly distinct pathways regulating synaptic autophagy in response to neuronal activity or metabolic paradigms.

### Synaptogyrin regulates degradation of synaptic vesicle components by autophagy

Newly produced synaptic vesicle proteins preferentially participate in neurotransmission and therefore the presynaptic terminals constantly turn over proteins to maintain neuronal function ^51^. Synaptic activity seems to control formation and likely function of autophagosomes and likely function ^16^ and prolonged neuronal firing can increase reactive oxygen species (ROS) production and oxidative damage of synaptic components ^52^. Moreover, although local translation takes place in a small number of axon terminals, the synthesis of synaptic vesicle-associated proteins is not sustained by this process ^53^. Therefore, the local turnover of synaptic vesicle proteins and vesicles themselves is critical for synaptic and neuronal function. Given that neuronal activity induces autophagosome formation at the synapse and that the synaptic vesicle protein Syngr regulates this process independently from the metabolically induced autophagy pathway, we asked whether autophagy mediates degradation of synaptic vesicle components via Syngr. Previous data show a reduction of synaptic vesicles in *syngr* null mutant NMJ synapses ^42^. To understand how Syngr couples synaptic autophagy with synaptic vesicle homeostasis, we performed analysis and quantification of the synaptic vesicle proteins. Cysteine-string protein (Csp2) is a synaptic vesicle protein present at the NMJ presynaptic compartment of control and Syngr mutant larvae **(Figure 3 A and B)**. Quantification of this protein at the presynapse shows that the protein levels of Csp2 are reduced in *syngr* null mutant synapses compared to controls **(Figure 3 D).** To further clarify if the reduction of Csp2 protein levels in *syngr* null mutant synapse is caused by an alteration in the regulation of autophagy, we blocked the initial steps of autophagosome formation. Atg17, in mammals called FIP200, is a component of the ULK complex and acts at the initial nucleation step ^54^. We confirm that Csp2 protein levels are sensitive to autophagy degradation, as *atg17 ^-/-^* and *atg9 ^-/-^* mutant synapses have more Csp2 protein levels than wildtype synapses **(Figure S3 A-A” and B-B”)**. Consistently, we found that blocking autophagy in *syngr* mutants (double knockouts *atg 17^-/-^, syngr ^-/-^*) restores Csp2 protein levels to those observed in control synapses **(Figure 3 A,C and D)**. We further confirmed that Csp2 protein levels do not change in response to autophagy induction via amino-acid deprivation **(Figure S3 C-C”)**. However, Csp2 protein levels are significantly reduced after 30 min induction of neuronal activity-induced autophagy using Nefiracetam treatment **(Figure S3 D-D”).** These results support that the reduction of Csp2 protein levels might be due to induction of neuronal activity-induced autophagy caused by loss of *syngr*. Our data further suggests that in *syngr* null mutants, synaptic vesicles – or at least synaptic vesicles components - might be taken up by autophagosomes for degradation. Hence, we further investigated if Csp2 colocalizes with the mature autophagosome marker Atg8^mCherry^. We used Pearson correlation to analyze the colocalization of Csp2 with Atg8^mCherry^ puncta (corresponding to the autophagosome). In *syngr* null mutant animals we observed a significant colocalization of the synaptic vesicle marker Csp2 and the autophagy marker **(Figure 3 G-G”,H)** compared to synapses of fed animals and animals after amino-acid starvation **(Figure 3 E-E”,F-F” and H)**. Additional analysis using the synaptic vesicle component Synaptotagmin (Syt) gave similar results **(Figure 3 I-L).** We confirm that Syt protein levels are also sensitive to autophagy degradation, as atg9 ^-/-^ mutant synapses have higher Syt protein levels than wildtype synapses **(Figure S3 E-E).** These results support a role for Syngr and synaptic autophagy in response to neuronal activity in the uptake of synaptic vesicle components (e.g. Csp2 and Syt) into autophagosomes, while this is not the case for the autophagy that occurs after amino-acid starvation. This further suggests that Syngr controls degradation of synaptic vesicles by synaptic autophagy in response to neuronal activity. Next, we were interested to uncover how autophagy is linked to synaptic vesicle recycling and degradation. The transmembrane autophagic protein Atg9 has been classically known for its function in autophagosome biogenesis and facilitates the expansion of the phagophore by mediating lipid transfer via Atg2 from donor membranes ^55–57^. Interestingly, Atg9 has not only been linked to synaptic autophagy but also to synaptic vesicle cycling, although its precise function here remains largely enigmatic ^27^. Moreover, Atg9 has also been found to be present on a subset of synaptic vesicles ^28^. Based on our previous results, we speculated that Syngr might interact with Atg9 to regulate autophagy. Hence, we performed immunoprecipitations in synaptosome-enriched preparations from adult *Drosophila* heads expressing endogenous levels of Atg9^GFP^ and probed against Syngr. We previously confirmed the expression of this genomic Atg9^GFP^ construct in the fly adult central nervous system **(Figure S4 A-B”)**. We detected Syngr in Atg9^GFP^ immunoprecipitates from synaptosomal enriched preparations but not from preparations that expressed only Mito-GFP (GFP with a target sequence for mitochondria) **(Figure 4 A).** Quantification of immunoprecipitation assays indicated a significant enrichment of Syngr in Atg9^GFP^ but not in Mito-GFP precipitates **(Figure 4 A’)** indicating that the binding of Syngr and Atg9 is specific. These results are in line with the work from Binottia *et al*. showing that immune-isolated Atg9 containing vesicles at the presynaptic terminal are specialized type of vesicles containing Syngr1 and Syng3 as well as members of the LC3 protein family and the selective autophagy receptor Sequestosome1 ^58^.

**Figure 3.**
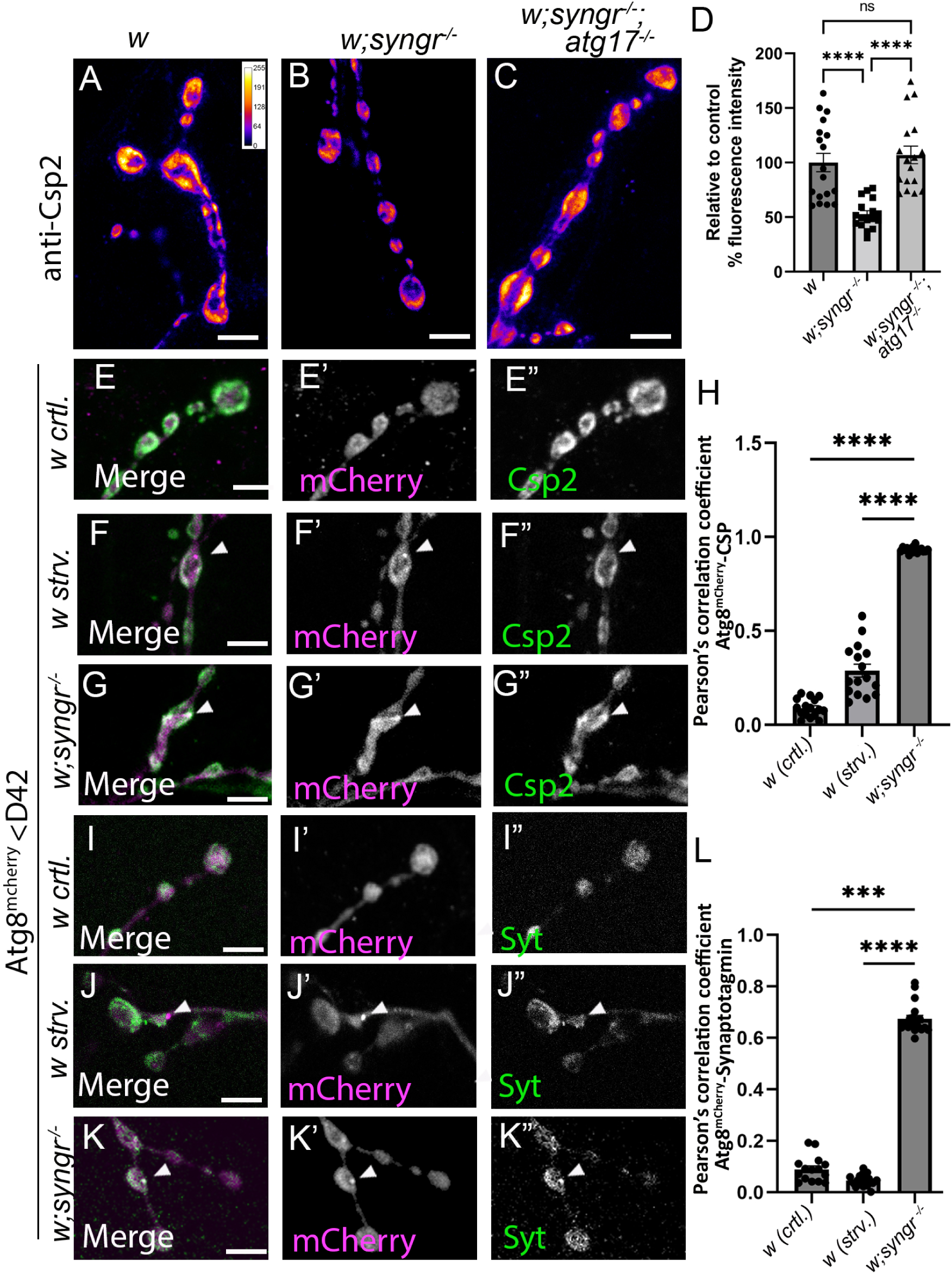
Synaptogyrin is necessary to control the degradation of synaptic vesicle components by neuronal activity-induced autophagy. (A-C) Immunostainings of synaptic vesicle component Csp2 in wildtype **(A)**, *syngr* null mutant (w;syngr^-/-^) **(B)** and *syngr-atg17* double null mutant (w;syngr^-/-^;atg17^-/-^) NMJs **(C)**. Fluorescence intensities are shown using the indicated scale in **(A)**. **(D)** Quantification of fluorescent intensities indicates a significant reduction of the synaptic vesicle marker Csp2 in *syngr* null mutants compared to wildtype and *syngr-atg17* double mutants. **(E-G”)** Immunostaining of wildtype control, 4h starved wildtype and fed *syngr* null mutant synapses. **(E,F).** Merge shows autophagosome marker Atg8^mCherry^ (magenta) **(E’,F’,G’)** and synaptic vesicle marker Csp2 staining (green) **(E’’,F”,G”)**. Csp2 staining is enriched in Atg8^mCherry^ positive dots (arrows) in *syngr* null mutant synapses (**G**, white). **(H)** Quantification of Pearson coefficient of Csp2 colocalization on Atg8^mCherry^ puncta in the mask of Atg8^mCherry^ puncta. **(I-K”)** Immunostaining of wildtype control, 4h starved wildtype and fed *syngr* null mutant synapses. **(I,J,K).** Merge shows autophagosome marker Atg8^mCherry^ (magenta) **(I’,J’,K’)** and synaptic vesicle marker Synaptotagmin (Syt) staining (green) **(I’’,J”,K”)**. Syt staining is enriched in Atg8^mCherry^ positive dots (arrows) in *syngr* null mutant synapses (**K**, white). **(L)** Quantification of Pearson coefficient of Syt colocalization on Atg8^mCherry^ puncta in the mask of Atg8^mCherry^ puncta. n >20 individual synapses from a minimum of 5 larvae. Scale bars= 5 µm.

**Figure 4.**
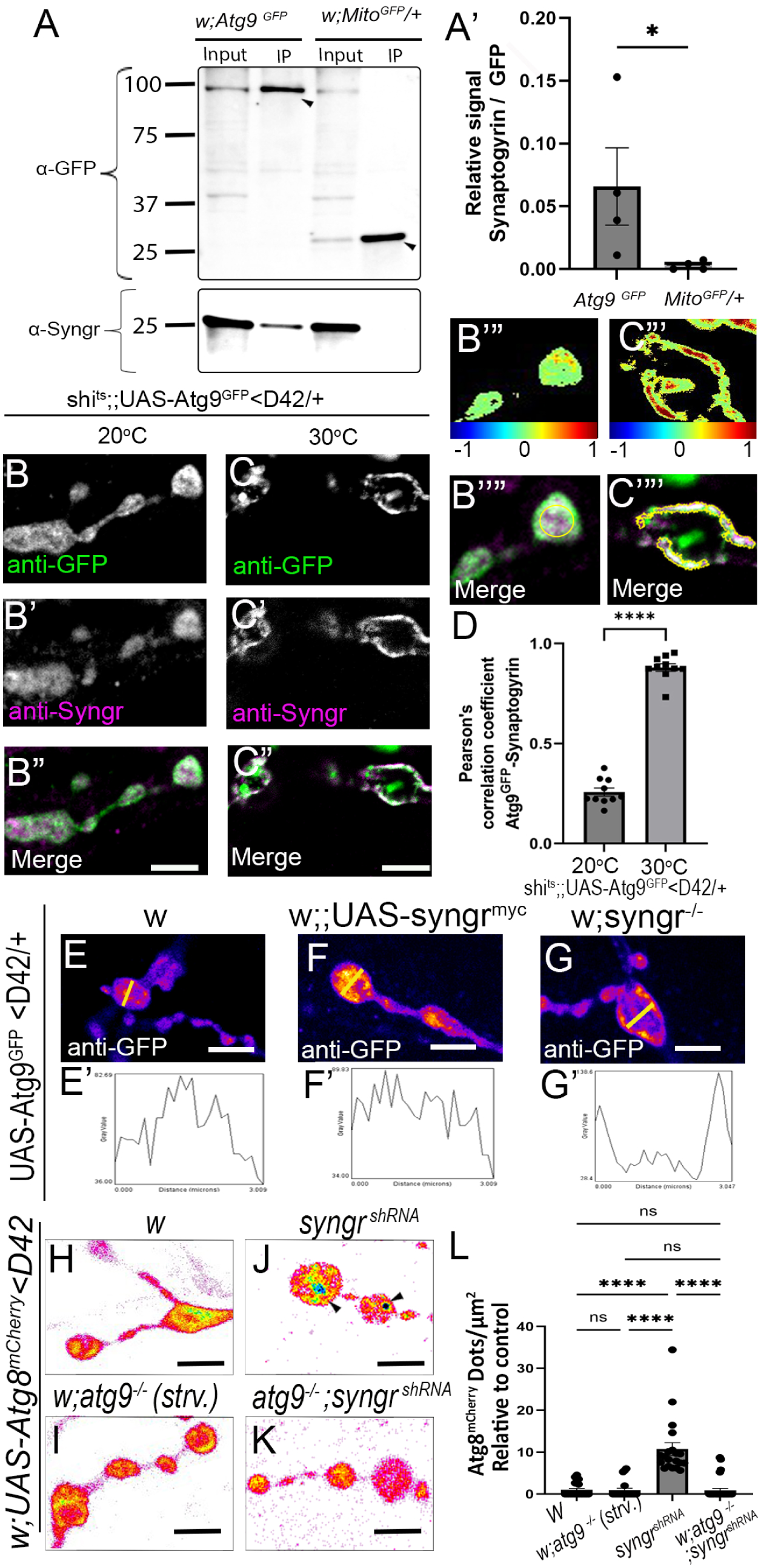
Synaptogyrin interacts with Atg9. **(A)** Western blot of immunoprecipitation assay (IP) using a GFP-Selector of synaptosomal-enriched preparations from fly heads expressing Atg9-GFP under endogenous expression levels and Mito-GFP (control). Blots were probed with anti-GFP (control Western Blot) to assess whether Syngr binds to Atg9. Note pulled-down GFP fusions are marked by arrowheads in the blot. **(A’)** Quantification of enrichment ratio of Syngr signal versus GFP signal. **(B-C”’)** Analysis of Atg9::GFP and Syngr immunostaining at larval NMJ in *shi^ts^*^1^ mutant background under non treated condition at 20°C **(B-B”’’)** and after depletion of synaptic vesicles by KCl stimulation at the non-permissive temperature of 30°C **(C-C”’’)**. **(B”’,C’”)** Colocalization colormaps of images B-B” and C-C”, respectively. Color code scale corresponds to normalized mean deviation product (nMDP). Values above 0 indicate colocalization. (**B’”’, C’’”)** Quantification mask in yellow define cytosolic area or synaptic vesicle pool close to the plasma membrane respectively. **(D)** Quantification of Pearson coefficient of Atg9-GFP colocalization with Syngr within defined yellow mask. **(E-G)** Immunostaining of Atg9^GFP^ in NMJ boutons of wildtype **(E)**, Synaptogyrin overexpressing (*syngr^myc^*) **(F)** and *synaptogyrin* null mutant flies (*syngr^-/-^)* **(G)** with corresponding distribution profile (along yellow line) (E’-G”). **(H-K)** Imaging of Atg8^mCherry^ puncta in NMJ boutons of wildtype **(H)**, 4h starved *atg9* null mutant flies (*w;atg9^-/-^*) **(I)**, flies expressing shRNAi against *syngr* **(J)** and flies expressing shRNAi against *syngr* in *atg9* null mutant background **(K).** Quantification of Atg8^mCherry^ puncta shows statistically significant reduction in *Syngr* knock down in *atg9* mutant background compared to wildtype background **(L).** n >20 individual synapses from a minimum of 5 larvae. Scale bars= 5 µm.

Given that Atg9-positive vesicles also undergo endocytosis at the presynaptic site ^27^, we tested whether Atg9 colocalizes with Syngr in these synaptic vesicles. To distinguish whether Atg9^GFP^ is localized to the synaptic vesicle pool or to the cytoplasm, we took advantage of the temperature sensitive *shibire^ts^* mutation (shi^ts^) (which encodes a dynamin orthologue) to sequester the synaptic vesicle pool located near the plasma membrane. The *shibire^ts^* mutation leads to a paralysis of endocytosis when animals are shifted from permissive temperatures (20°C) to a higher, non-permissive temperature of 30°C. Subsequent neuronal stimulation leads to fusion of synaptic vesicles with the plasma membrane, leaving within the presynapse only those synaptic vesicles that are not ready for neurotransmitter release. Consequently, we found that at 20°C Atg9^GFP^ and Syngr are widely distributed at the *Drosophila* NMJ synapse **(Figure 4 B-B’’)**, but when these animals are shifted to 30°C followed by potassium chloride (KCl) stimulation, both Syngr and Atg9^GFP^ colocalize nearby the plasma membrane **(Figure 4 C-C”)**. We analysed colocalisation by creating a colocalisation map that allows a quantitative visualisation of colocalisation and show high colocalisation levels (represented as hot colors) in some areas of the bouton between Atg9-GFP and Syngr when animals are shifted to 30°C followed by KCl stimulation (**Figure 4 B’’’,C’’’)**. Next we quantified the colocalization between Atg9 and Syngr in these two conditions using Pearson’s correlation coefficient (**Figure 4 B””,C”” and D**). Changes in synaptic Atg9 localization occur during synaptic autophagy in response to neuronal activity and mislocalization of this autophagy protein is associated with defects in neuronal activity-induced synaptic autophagy ^27^. Consistently, we observed than overexpression or deletion of Syngr (that in turn represses or activates synaptic autophagy, respectively) lead to changes in the distribution of Atg9 at the synapse **(Figure 4 E-G)**. Next, we tested if Atg9 and Syngr interact to mediate autophagy. First we confirmed that synaptic autophagy is blocked in *atg9* null (*atg9^-/-^*) mutants **(Figure 4 H,I and L)** and found that the increase in autophagy caused by knockdown of Syngr (*syngr^shRN^*^A^) was significantly decreased in *atg9^-/-^* mutant flies (*syngr^shRNA^; atg9^-/-^)* **(Figure 4 J-L)** indicating that *atg9* is epistatic over *syng.* Moreover, *atg9^-/-^* mutants that abolish autophagy show higher levels of Syngr within the synaptic vesicle pool compared to *atg9^-/-^* mutant synapses expressing a genomic *atg9* rescue construct (*atg9^-/-^;* atg9^HA^) **(Figure S4 C-E)**. These data are supported by the fact that Syngr has been shown to be a target of basal autophagy ^59^.Taken together our results support the link between Atg9 and synaptic vesicle cycle via Syngr, and the degradation of synaptic vesicle components via neuronal activity-induced autophagy.

### Synaptogyrin ameliorates synaptic autophagy defects present in a *Drosophila* model of Frontotemporal dementia

Next we wanted to test the relevance of Syngr in neurological disease since pathological Tau binds to mammalian Syngr-3 and studies in *Drosophila* and vertebrate models suggests that this binding mediates the accumulation of pathological Tau on synaptic vesicles ^32^. Given this relationship we wondered if pathological Tau also alters synaptic autophagy. Therefore, we expressed human Tau with a pathological mutation for Frontotemporal dementia (FTD) (hTau^R406W^) in motor neurons and observed that this leads to an accumulation in the number of autophagic vesicles similar to the increase observed upon autophagy stimulation (**Figure 5 A-C).** This suggests that expression of this pathological hTau^R406W^ might lead to an increase in the biogenesis or in the accumulation of autophagosomes. However, mutations in Tau have been associated to alterations in axonal transport ^60^ and to impairment of autophagy flux ^61^. Thus, we further analyzed portion of autophagosomes and autolysososmes using the Atg8mCherry-GFP probe and observed that expression of hTau^R406W^ leads to significantly higher quantities of mCherry (autolysosomes) than mCherry-GFP (autophagosome) fluorescent Atg8-positive puncta compared to control synapses where autophagy is induced **(Figure 5 D,E,G and S5 A,B)**. This indicates that expression of the pathological Tau protein leads to the increased amount of autolysosomes at the presynapse. Intriguing, it has been shown that mice harboring a single copy of the *syngr-3* gene showed an improvement of the Tau-dependent memory defects ^38^. We therefore asked if reducing the levels of Syngr could have a positive effect on autophagy alterations caused by hTau^R406W^. Indeed, our results showed that removing one gene copy of *syngr* (*syngr^+/-^*) not only activates autophagy (consistently with our previous data showing this effect when *syngr* is knocked down) but also rescues the autophagic alterations quantified in the NMJ synapses from animals expressing hTau^R406W^ **(Figure 5 F-G and S5C)**. Evidence supports that Tau is degraded, at least partially, by macroautophagy ^62^. Accordingly, reducing the levels of Syngr to activate synaptic autophagy results in a reduction of the levels of pathological hTau^R406W^ at the synapse, while inhibition of autophagy in these animals restores synaptic Tau levels similar to control (**Figure 5 H-K**). To strengthen these results we performed colocalization studies and confirmed that activating synaptic autophagy by reducing the levels of Syngr significantly increases the colocalization between of Tau and the autophagosomal marker Atg8, further indicating that induction of synaptic autophagy by reducing the levels of Syngr favors the recruitment of Tau into the autophagosome at the synapse (**Figure 5 L-N**). Finally, we asked whether the degradation of pathological Tau was specific for the loss of Syngr. Intriguingly, induction of autophagy by amino-acid starvation does not result in lower levels of pathological Tau at the synapse (**Figure 5 O-Q**), suggesting that activation of autophagy by loss of Syngr have distinct physiopathological roles at the synapse. Our data suggests that targeting Syngr stabilization and degradation could result in novel therapeutic approaches to exploit specifically neuronal activity-induced autophagy to treat FTD and likely other neurodegenerative-related diseases.

**Figure 5.**
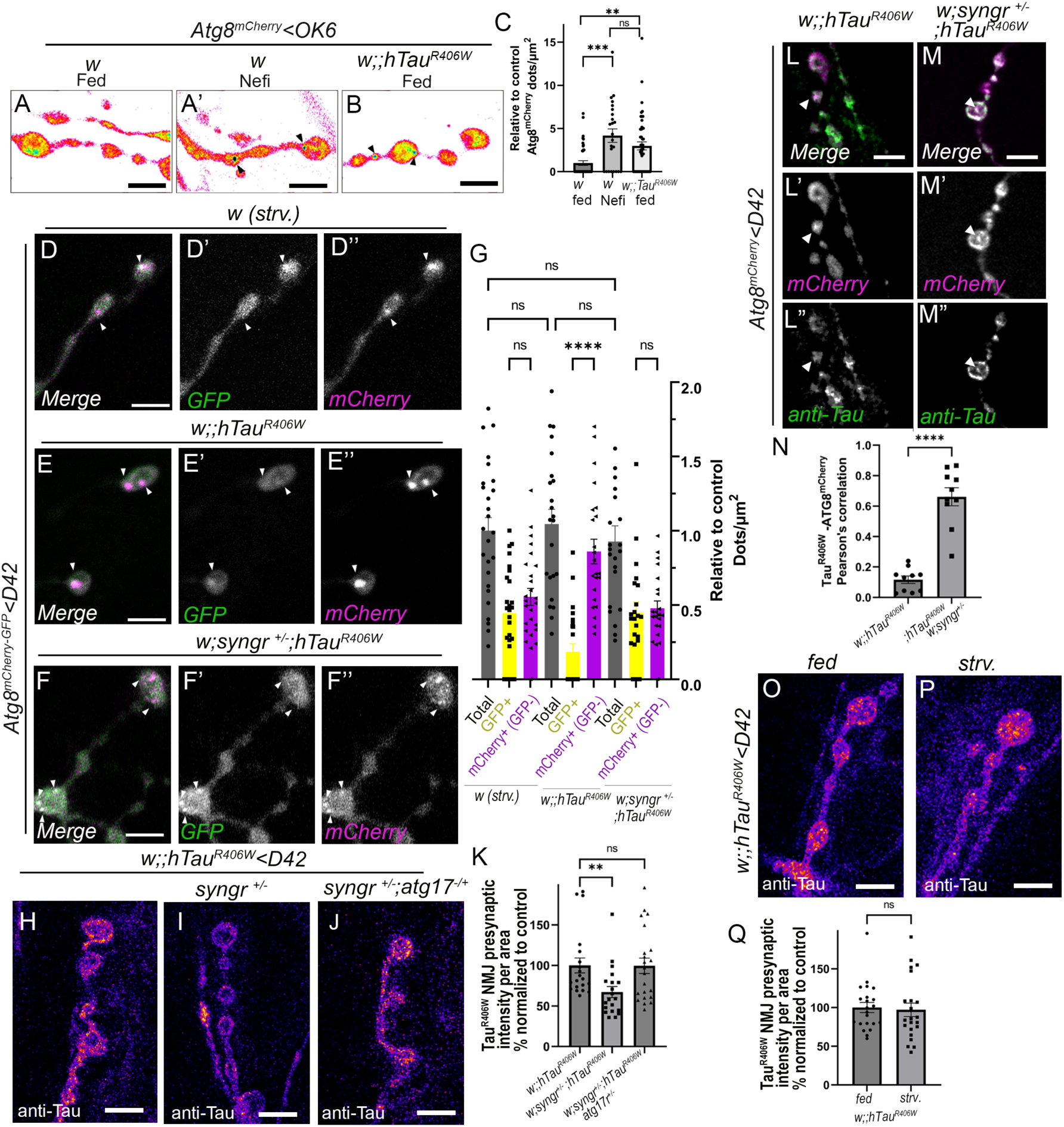
Synaptogyrin participates in the degradation of pathological Tau via autophagy. (A-B) Imaging of Atg8^mcherry^ puncta formation at the synapse of indicated genotypes. **(A)** wildtype fed, wildtype after neuronal stimulation with Nefiracetam (Nefi) showing autophagosomes (arrowheads) **(A’),** and human hTau^R406W^ (pathological mutation) expressing synapses showing autophagosomes (arrowheads) **(B)**. Quantification shows statistically significant increased Atg8^mcherry^ puncta formation after neuronal stimulation and in synapses expressing hTau^R406W^ **(C).** n >20 individual synapses from a minimum of 5 larvae. **(D-F”)** Live imaging of Atg8^mCherry-GFP^ to analyze amount of autophagosomes and autolysosomes in NMJ boutons of flies after 4h starvation **(D-D”),** flies expressing hTau^R406W^ **(E-E”),** and of flies expressing hTau^R406W^ with reduced levels of Syngr (*syngr* ^-/+^) **(F-F”)**. **(G)** Quantifications of the total number of Atg8 dots (GFP+mCherry plus mCherry alone) and the number of white (GFP+mCherry) or magenta (mCherry) Atg8 dots per NMJ area. Note that hTau^R406W^ expressing synapses have significantly lower Atg8 GFP positive autophagosomes (before acidification by fusion with lysosomes) compared to wildtype animals with induced autophagy or hTau^R406W^ with reduced levels of Syngr. N >27 individual synapses from a minimum of 5 larvae. **(H-J)** Immunostaining of anti-Tau in synapses expressing hTau^R406W^ **(H),** in synapses expressing hTau^R406W^ with reduced levels of Syngr (*syngr* ^-/+^) **(I),** and in synapses expressing hTau^R406W^ with reduced levels of Syngr and Atg17 (*syngr* ^-/+^; atg17^-/-^) **(J).** Quantification shows a statistically significant reduction of Tau staining when Syngr levels are lowered and this reduction is not observed in in animals with lower Syngr and Atg17 levels **(K).** n >20 individual synapses from a minimum of 5 larvae. **(L-M”)** Tau and Atg8^mCherry^ immunostaining of synapses expressing hTau^R406W^ in wildtype background **(L)** and with lower levels of Syngr **(M).** Merge shows autophagosome marker Atg8^mCherry^ (magenta) **(L’,M’)** and Tau (green) **(L’’,M”)**. Tau staining is enriched in Atg8^mCherry^ positive dots (arrows) in synapses with lower levels of Syngr (**M**, white). **(N)** Quantification of Pearson coefficient of Tau colocalization on Atg8^mCherry^ puncta in the mask of Atg8^mCherry^ puncta. n >18 individual synapses from a minimum of 5 larvae. Tau immunostaining in fed and 4h starved animals expressing hTau^R406W^ **(O,P)**. Quantification of fluorescent intensities shows no significant difference between fed and 4h starved animals **(Q).** n >24 individual synapses from a minimum of 5 larvae. Scale bars= 5 µm.

## Discussion

To fully understand how autophagy participates in the nervous system at cellular level, it is essential to gain valuable insights concerning brain function during health and disease. However, our current understanding of how autophagy contributes to neuronal function and survival is far from complete. Moreover, how autophagy is regulated in different cellular compartments remains enigmatic. This is especially important in neurons that are highly polarized and specialized post-mitotic cells where different compartments exert distinct roles during neuronal function and communication. Indeed, previous studies reported the existence of differences in autophagosome dynamics in dendrites and axons ^16^. Moreover, synaptic proteins with previously known roles in synaptic vesicle formation and recycling such as Endophilin A or Synaptojanin and the active zone protein Bassoon have been described to play additional roles in synaptic autophagy ^10,12,14,15,30^. Besides this evidence, whether synaptic autophagy undergoes distinct molecular adaptations and functions in response to neuronal activity or metabolic stimuli is mostly unknown. Here we provide novel data showing that neuronal activity and metabolic signals operate via distinct molecular mechanisms to regulate autophagy at the neuronal synapse, and are likely to support different aspects of cellular homeostasis. Syngr is a highly conserved, abundant structural protein of synaptic vesicles ^28^ and loss of Syngr has been shown to alter synaptic vesicle size and activity of membrane transporters ^42,63^. However, the precise functions of Syngr are still not well described. Here we report a novel function for this protein in the regulation of synaptic autophagy in response to neuronal activity (see graphical summary **Figure S5 D)**. We show that loss of Syngr leads to constitutive autophagy induction without affecting the autophagy flux (fusion of the autophagosome with the endolysosome to degrade autophagic cargos). Consistently, we show that overexpression of Syngr blocks autophagy induction in response to neuronal stimulation. These results indicate that Syngr is a negative regulator of autophagy. Previous work has described how mutations in the synaptic proteins Endophilin-A or Synaptojanin block autophagosomal formation after both neuronal activity and amino-acid starvation ^10,12,14^. In contrast, we report that overexpression of Syngr has no effect in autophagy induction after amino-acid deprivation, describing for the first time the role of a protein that participates solely in the autophagy pathway activated by neuronal activity. We believe this work will allow additional studies to fully decipher how neuronal activity regulates synaptic autophagy in health and disease. We recently published that EndoB, a presynaptic-enriched protein, is required for autophagy induction after amino-acid deprivation and during photoreceptor development ^39^. Here we now show that Syngr and EndoB function independently from each other since loss of EndoB does not block increased autophagosome formation in animals where Syngr is knocked down. Thus, our results confirm the existence of two pathways regulating autophagy at the synapse: one metabolic that responds to amino-acid deprivation via EndoB, and a second one that is triggered by neuronal activity through Syngr. Recent research proposes that autophagy might turnover synaptic vesicles particularly during intense neuronal activity ^28,51^. This idea stems from the fact that a small number of autophagic proteins have been found on synaptic vesicles, but also that during intense neuronal activity synaptic vesicles are reused many times and that this might lead to less fusion-competent synaptic vesicles over time ^28,51^ and consequently affects fidelity during neurotransmission. Therefore, it seems interesting to think that autophagy might help to strengthen neurotransmission by turning over synaptic vesicles. In fact, neuronal knockdown of Atg7, a protein of the autophagic core machinery, decreases neurotransmission ^64^, thus strengthening the notion that autophagy at presynaptic site is required to maintain functional neuronal communication. However, how the processes of autophagy, neuronal activity, and synaptic vesicle turnover are linked to each other still remains poorly understood. Here we found evidence indicating that neuronal activity-induced autophagy plays a role in the regulation of synaptic components. We show decreased levels of Csp2, a marker of synaptic vesicles, in *syngr* null mutant presynaptic terminals. These results are in line with previous observations that showed a decreased number of synaptic vesicles in *syngr* null mutants ^42^. We here show that blocking the initial steps of autophagy re-establishes Csp2 levels in *syngr* null mutant synapses, indicating that the reduction of Csp2 quantified in *syngr* null mutants is, at least in part, caused by an increase in autophagy. Moreover, we show that induction of neuronal activity-induced autophagy but not starvation-induced autophagy results in lower levels of Csp2. Our data further suggest that synaptic vesicle components (e.g. Csp2 and Syt) are taken up into autophagosomes in *syngr* null mutants and that this is not the case for autophagosomes that are formed after amino-acid starvation. Hence our results suggests that Syngr regulates synaptic vesicles turnover via neuronal activity-induced autophagy. Interestingly, the autophagic core protein Atg9 has been found on some synaptic vesicles ^28^, being proposed as a link between autophagy and synaptic vesicle cycle ^27^ and likely also in the degradation of synaptic vesicles. Here we provide the missing link and show that Syngr binds to Atg9 and that these two proteins colocalize in presynaptic terminals and that Atg9 distribution is dependent on Syngr levels, likely through Syngr-Atg9 binding. Furthermore, we show that knock down of Syngr in *atg9* null mutants does not lead to an increase in autophagy, indicating that both Atg9 and Syngr functions are required to regulate autophagy induction at presynaptic terminals.

Interestingly, reduction of Syngr can rescue pathological traits in a mouse model of FTD expressing a pathological form of Tau ^32,38^. However, the cellular process that mediates this rescue is unclear. We now provide evidence that expression of pathological Tau leads to the accumulation of autolysosomes by impairing autophagy and that a reduction of Syngr levels can rescue this phenotype by restoring appropriate autophagy function. Moreover, we show that reduction of Syngr levels reduce the amount of pathological Tau in presynaptic terminals and that this is dependent on autophagy. Moreover, we show that pathological Tau co-localizes with Atg8-positive autophagosomes in animals with lower levels of Syngr, indicating that Syngr regulates the degradation of pathological Tau via autophagy. By contrast, in animals with normal levels of Syngr pathological Tau does not significantly colocalize with Atg8 positive autophagosomes indicating that Tau degradation via autophagy is regulated by Syngr levels. Furthermore, our data point out that amino-acid deprivation induced autophagy is not actively degrading pathological Tau, sustaining the idea that different stimulus (amino -cid deprivation and neuronal activity) function over different molecular autophagy pathways at the presynaptic terminal to execute different functions. Taken together our results indicate that activation of autophagy by lowering Syngr levels mediated degradation of pathological Tau via the neuronal activity dependent autophagy pathway. However, further research is required to understand the precise molecular mechanism of Syngr-mediated Tau degradation.

Our work strengthens the knowledge about the regulation of autophagy in neurons in health, but it also could open new avenues into exploiting autophagy in neurodegenerative diseases. We describe for the first time how the synaptic protein Syngr functions in autophagy in response to neuronal activity and also provide additional mechanistic data about the role of Syngr and neuronal autophagy in FTD. In summary, our work reveals a novel molecular mechanism regulating the quality control of synaptic vesicles, a key step in maintaining the fidelity of neurotransmission and synaptic decay. Moreover, the identification of synaptic proteins that solely participate in neuronal-induced autophagy could lead to therapeutic strategies to specifically target neuronal autophagy in the context of neurodegenerative diseases.

## Materials and Methods

### Fly Stocks, genetics and detailed genotype description

Flies were grown on standard cornmeal and molasses medium supplemented with yeast paste at 25°C. The following *Drosophila* stocks were obtained from the Bloomington Stock Center: *w*; P{w[+mW.hs]=GawB}D42, P{w[+mW.hs]=GawB}OK6, w* P{GSV1}C155^C155^*, w;UAS-eGFP, *w*; P{UAS-TrpA1(B).K}attP16, w* P[UAS-TrpA1(B (B).K}attP2.,* y^1^ sc* v^1^ sev^21^; P{TRiP.HMS01724}attP40 (also called UAS-syngr^shRNA^) and shi^ts1^. The following fly stocks were obtained from the lab where the stock was first described: *w; endoB*^4^ was described in ^39^, w;;UAS-Atg18::GFP and *w;; UAS- mCherry-GFP::Atg8 in* ^12^, w;;*UAS-GFP::Atg9* in ^10^, *w; GFP::Atg9* (P.Verstreken, unpublished), *w; P[w{+}UAS-Atg8-mCherry* in ^63^, w;;UAS-hTau ^R406W^ in ^64^, *w; P[w{+}UAS-Atg8-mCherry and w;P[w{+}UAS-Atg8-mCherry* in ^63^, *w; syngr* ^1^ *and w;; UAS- syngr::myc* in ^42^*, w; atg9^9B5^; HA::atg9* in ^65^ *and w;;atg17^d^*^130^ ^67^.

Stocks were grown on standard cornmeal/agar medium at 25°C.

For a full list of used genotypes see supplementary information.

## Materials and Methods

### Autophagy assay

Autophagy experiments were carried out as previously published ^10^. For amino acid starvation experiments flies were raised at low population density on standard fly food supplemented with yeast paste at 25°C. Early third instar larvae were placed on Petri dishes containing 20% sucrose and 1% agarose for 3–4 hr prior to dissection.

For autophagy induction via Rapamycin, third-instar larvae were fed for 4 h on standard food and standard food containing 5μM Rapamycin (Sigma).

For autophagy induction by neuronal stimulation, TrpA1 (D42-Gal4)-expressing larvae grown at 20°C were placed in a preheated 30°C tube for 30 min and dissected in heated HL3.

Pharmacological autophagy induction using calcium channel agonist treatment was performed as previously described with minor modifications ^14^. In brief, larvae were dissected on sylgard plate in HL3 solution and incubated for 30min in Nefiracetam solution (1 μM Nefiracetam (Tocris) in 0.0025% DMSO and 1 mM Ca2Cl in HL3) or control solution (0.0025% DMSO and 1 mM Ca_2_Cl in HL3). Animals were subsequently washed 3 time with HL3 and fixed as usual.

For calcium chaletor treatment larvae were dissected on syrgard plates in HL3 and incubated for 30 min in calcium chaletor solution (25 µM.EGTA-AM (Thermo Fisher), 1 mM CaCl_2,_ 0.0025% DMSO in HL3) or control solution (1 mM CaCl_2_ and 0.0025% DMSO in HL3)

### Synaptic vesicle depletion assay

Third-instar larvae were dissected in HL3 buffer without Ca2^+^ on a sylgard plates at room temperature. Following the dissection, the HL3 buffer was exchanged with a 34 °C warm Ca2^+^-free HL3 buffer and immediately placed in an oven at 34 °C for 2 min to adjust the preparation temperature. The preparation was stimulated with preheated HL3 supplemented with 90 mM KCl and 1.5 mM Ca2^+^ for 5 min at 34 °C. Following stimulation, the preparation was incubated two times for 3 min each with Ca2^+^-free HL3 at 34 °C. The samples were then fixed in pre-warmed 4% PFA at 34 °C for 15 min and processed for immunohistochemistry.

### Immunohistochemistry

Immunohistochemistry of cryosections from *Drosophila* brains was done following a standard procedure. 1-3 days adult fly heads were dissected and fixed in 4% PFA and HL3 for 40 min at RT. The heads were washed in PBS and cryo-preserved in 10% sucrose for 30 min, followed by overnight incubation in 30% sucrose. Heads were immersed in tissue-freezing medium (Tissue-Tek OCT), frozen on dry ice and 10μm-thick cryosections were collected on Superfrost slides. Sections were permeabilized in 0.1% Triton X- 100 in PBS (PBT).

For immunohistochemistry of pupal and adult *Drosophila* brains, animals were dissected in ice-cold PBS, transferred into tubes and fixed with 4% in PBS for 45 min at RT.

Immunohistochemistry of third instar larval NMJs were performed by dissecting larvae in HL3 (110 mM NaCl, 5 mM KCl, 10 mM NaHCO3, 5 mM Hepes, 30 mM sucrose, 5 mM trehalose, and 10 mM MgCl2, pH 7.2) on sylgard plates. Dissected *Drosophila* larvae were fixed with paraformaldehyde 4% in HL3 for 20 min at RT, transferred into tubes and permeabilized for 45 min with 0.4% Triton in PBS.

Samples from cryosections of adult brains and third instar larvae were pre-blocked with 10% NGS-PBT (0.1% Triton and 0.4% Triton in PBS respectively) for 30-60 min at RT. Samples were incubated with the primary and secondary antibodies in PBT-10% NGS overnight at 4°C and for 2 hr at RT, respectively. For imaging, samples were mounted on Vectashield (Vector Laboratories).

The following primary antibodies were used: chicken α-GFP [1:1000 (Invitrogen; A10262)], chicken α-GFP [1:1000 (AVES; 1020), chicken α-mCherry [1:1000 (Novus; NBP2-25158)], mouse α-Dlg-PDZ 4F3 [1:500 (Developmental Studies Hybridoma Bank, DSHB)], mouse α-csp2 (6D6) [1:100 (DSHB)], mouse α-Syt 3H2 2D7 [1:200 (DSHB)], mouse α-hTau (HT7) [1:4000 (Thermo Fisher)] and rabbit α-Syngr [1:500 ^42^ gifted by J.T .Littleton]. Alexa Fluor 488-/Alexa Fluor 568-conjugated secondary antibodies were used at 1:1000 (Invitrogen).

### Imaging, image acquisition and quantitative image analysis

For imaging of autophagic markers at the NMJ, third instar larvae were washed in HL3, dissected in fresh HL3 and pinned on Sylgard plates. For live imaging, animals were immediately imaged in fresh HL3 with the HCX APO L 63x 0.90 W U-V-I water dipping lens on a SP5-MP (Leica Microsystems) confocal microscope. Fixed brains were imaged using a Leica HCX PL APO lambda blue 40×/1.25 OIL UV or HCX PL APO lambda blue 63×/1.40-0.60 OIL UV objective on a TCS SP5 (Leica Microsystems) and PL APO 63x/1.40 Oil CS2 objective on a Leica Stellaris 5 confocal microscope.

Samples from 5 larvae per genotype/condition were imaged through single confocal planes, imaging 4 NMJs per larvae. To guarantee representability, each NMJ image was taken from a different segment (A2, A3 or A4 corresponding to the muscles 12 and 13) and two from each side (left-right) of the larvae. Brains were scanned by confocal microscopy under identical settings. Quantification of number of puncta or intensity was performed in NIH ImageJ. Number and size of autophagosomes (Atg8-mCherry positive puncta) were quantified using the previously published automatic quantification method “Autophagoquant” ^43^.

All images were processed using ImageJ and figures were composed in Adobe Photoshop 2021. Fluorescence intensities are color-coded using Lookup tables with inverted 16 and fire colors.

Colocalization of Atg8mCherry dots with Csp2 and Syt was calculated using JACoP in ImageJ, where an object-based Pearson correlation coefficient is defined by thresholding the Atg8mCherry puncta (thresholded overlay colocalization coefficient). Colocalization Colormap of Atg9-GFP and Syngr was performed according to ^70^ using a Colocalization Colormap ImageJ plugin.

### Immunoprecipitation (IP)

Adult flies with the following genotypes (w, *UAS-Mito-eGFP/+* and *w; GFP::Atg9)* were frozen on dry ice and decapitated by vigorously vortexing. Heads were separated from the body by sieving. 22 mg of heads for each genotype were homogenized on ice using a motorized pestle in 1 ml of homogenization buffer containing 300 mM sucrose, 400 mM HEPES (pH7.4), protease inhibitor cocktail cOmplete^TM^ EDTA-free and PhosStop^TM^ (Roche). The homogenate was cleared from debris by centrifugation for 10 min at 1000 x g twice and NP40 to a final concentration of 0.5% and Sodium Chloride to a final concentration of 150 mM were added to the supernatant. Samples were incubated on ice for 20 minutes and 60 μl of the supernatant were kept for posterior analysis and the rest of supernatant was loaded onto 30 μl of GFP Selector Magnetic beads (NanoTag Biotechnologies) and incubated for 1 hour at 4°C on a shaker. GFP-Selector was magnetically separated and washed three times with 10 mM HEPES pH7.5, 150 mM NaCl, 0,5% NP40 and protease inhibitor cocktail cOmplete^TM^ EDTA free and PhosStop^TM^. For the gel analysis, beads were resuspended in 30 μl PBS and 10 μl 4xSDS buffer (Bio Rad), denatured for 15 min at 75°C and separated by SDS-PAGE on a 4-20% gradient Gel. For Western blots, SDS-PAGE gels were blotted onto PDVF transfer membrane (Millipore) and incubated with antibodies using standard procedures. Antibody dilutions used: α-GFP (Aves Labs) 1:1000, anti-Syngr 1 ^42^, peroxidase-conjugated secondary antibodies against rabbit 1:5000 (Invitrogen) and against chicken 1:1000 (Proteintech). The ECL system (Perkin Elmer) was used for detection and chemiluminescence was imaged using ChemiDoc™ Touch Imaging System (Bio Rad). Quantification of western blot data was performed using Image J software.

### Statistics

Statistical significance was determined with one-way analysis of variance tests (ANOVA), followed by Turkey-Kramer testing using GraphPad Prism software, Version 9 (Graphpad). The distribution of data was analyzed using a D’Agostino-Pearson Omnibus test. Normal data were tested with parametric tests: for two datasets we used a student’s t-test. For more than two datasets, we used a one-way analysis of variance test (ANOVA) and a post hoc Tukey test. Non-normal distributed data were tested using non-parametric tests: for two datasets we used a Mann-Whitney test and for more than two datasets an ANOVA-Kruskal Wallis test and a Dunn’s test were used. Significance of statistical difference was defined as ****=p<0.0001, ***=p<0.001, **=p<0.01, *=p<0.05, ns= p>0.05. The number of analyzed samples (NMJ boutons, brains or laminas) is indicated in the figure legend.

## Acknowledgments

We thank Jesus Diaz Cuevas for technical support of the experiments, T.Littelton, P.Verstreken, Neufeld, G. Juhasz, the Bloomington Drosophila Stock Center, FlyORF and the Developmental Studies Hybridoma bank for reagents. We thank Prof. Volker Haucke (Leibniz Institute for Pharmacology), Prof. Eckart D. Gundelfinger (Leibniz Institute for Neurobiology), Dr. Benjamin Dehay, and Dr. Erwan Bezard (IMN/CNRS/University Bordeaux) for critical reading and valuable comments on the manuscript.

SFS received financial support from the PID2023-153022NB-100 project financed by MCIN/AEI/10.13039/501100011033 /FEDER, EU, Ikerbasque Foundation for Science, Initiative d’Excellence de l’Université de Bordeaux (IDEX Neurocampus Chair), GPR BRAIN_2030 and the Region Nouvelle-Aquitaine. SFS acknowledge support from Marie Sklodowska-Curie COFUND Programme of the European Commission HORIZON-MSCA-2022-COFUND-101126600-SmartBRAIN3.

RB and JDS acknowledge support from the European Cooperation for Science & Technology ProteoCure COST Action (CA20113). RB and JDS were funded by MCIN/AEI/10.13039/501100011033, projects PID2023-147399NB-I00 and PID2020-114178GB-I00 and CEX2021-001136-S and CEX2021-001202-M Severo Ochoa Excellence Program and additional support provided by the Diputación Foral de Bizkaia, Programa Transferencia Tecnológica 2023 and the Department of Industry, Tourism, and Trade of the Basque Country Government (Elkartek Research Programs)

## Author Contributions

SFS conceived the study, SFS, SH-D, and PM-O analyzed data, SFS and SH-D wrote the paper. SH-D, PM-O, IS-M, CM, SG, JDC, JDS, RB, and SFS, performed experiments and/or provided reagents.

## Competing Interest Statement

The authors declare that the research was conducted without any commercial or financial relationships that could be construed as a potential conflict of interest.

## Notes

### Competing Interest Statement

The authors have declared no competing interest.

### Summary of Updates

Figure 3, 4 and 5 revised; author affiliations updated; Supplemental files updated.

## References

1. Mizushima N. The pleiotropic role of autophagy: from protein metabolism to bactericide. Cell Death Differ 2005; 12:1535–41.

2. Galluzzi L, Baehrecke EH, Ballabio A, Boya P, Bravo-San Pedro JM, Cecconi F, Choi AM, Chu CT, Codogno P, Colombo MI, et al. Molecular definitions of autophagy and related processes. EMBO J 2017; 36:1811–36.

3. Fleming A, Rubinsztein DC. Autophagy in Neuronal Development and Plasticity. Trends Neurosci 2020; 43:767–79.

4. Ariosa AR, Klionsky DJ. Autophagy core machinery: overcoming spatial barriers in neurons. J Mol Med (Berl) 2016; 94:1217–27.

5. Mortimore GE, Schworer CM. Induction of autophagy by amino-acid deprivation in perfused rat liver. Nature 1977; 270:174–6.

6. Mizushima N, Ohsumi Y, Yoshimori T. Autophagosome formation in mammalian cells. Cell Struct Funct 2002; 27:421–9.

7. Klionsky DJ, Abdel-Aziz AK, Abdelfatah S, Abdellatif M, Abdoli A, Abel S, Abeliovich H, Abildgaard MH, Abudu YP, Acevedo-Arozena A, et al. Guidelines for the use and interpretation of assays for monitoring autophagy (4th edition). Autophagy 2021; :1–382.

8. Komatsu M, Ueno T, Waguri S, Uchiyama Y, Kominami E, Tanaka K. Constitutive autophagy: vital role in clearance of unfavorable proteins in neurons. Cell Death Differ 2007; 14:887–94.

9. Maday S, Holzbaur ELF. Compartment-specific regulation of autophagy in primary neurons. J Neurosci 2016; 36:5933–45.

10. Soukup S-F, Kuenen S, Vanhauwaert R, Manetsberger J, Hernández-Díaz S, Swerts J, Schoovaerts N, Vilain S, Gounko N V., Vints K, et al. A LRRK2-Dependent EndophilinA Phosphoswitch Is Critical for Macroautophagy at Presynaptic Terminals. Neuron 2016; 92:829–44.

11. Kuijpers M, Kochlamazashvili G, Stumpf A, Puchkov D, Swaminathan A, Lucht MT, Krause E, Maritzen T, Schmitz D, Haucke V. Neuronal Autophagy Regulates Presynaptic Neurotransmission by Controlling the Axonal Endoplasmic Reticulum. Neuron 2021; 109:299–313.e9.

12. Vanhauwaert R, Kuenen S, Masius R, Bademosi A, Manetsberger J, Schoovaerts N, Bounti L, Gontcharenko S, Swerts J, Vilain S, et al. The SAC1 domain in synaptojanin is required for autophagosome maturation at presynaptic terminals. EMBO J [Internet] 2017; 36:1392–411. Available from: https://www.embopress.org/doi/10.15252/embj.201695773

13. Hill SE, Kauffman KJ, Krout M, Richmond JE, Melia TJ, Colón-Ramos DA. Maturation and Clearance of Autophagosomes in Neurons Depends on a Specific Cysteine Protease Isoform, ATG-4.2. Dev Cell 2019; 49:251–266.e8.

14. Bademosi AT, Decet M, Kuenen S, Calatayud C, Swerts J, Gallego SF, Schoovaerts N, Karamanou S, Louros N, Martin E, et al. EndophilinA-dependent coupling between activity-induced calcium influx and synaptic autophagy is disrupted by a Parkinson-risk mutation. Neuron 2023; 111.

15. Decet M, Soukup S-F. Endophilin-A/SH3GL2 calcium switch for synaptic autophagy induction is impaired by a Parkinson’s risk variant. Autophagy 2023; :1–3.

16. Kulkarni VV, Anand A, Herr JB, Miranda C, Vogel MC, Maday S. Synaptic activity controls autophagic vacuole motility and function in dendrites. J Cell Biol 2021; 220.

17. Kabeya Y, Mizushima N, Ueno T, Yamamoto A, Kirisako T, Noda T, Kominami E, Ohsumi Y, Yoshimori T. LC3, a mammalian homologue of yeast Apg8p, is localized in autophagosome membranes after processing. EMBO J 2000; 19:5720–8.

18. Malinowska M, Kloska A, Magdalena W, Gabig-cimi M. Lipophagy and Lipolysis Status in Lipid Storage and Lipid Metabolism Diseases. 2020; 1:1–31.

19. Kim D-H, Sarbassov DD, Ali SM, King JE, Latek RR, Erdjument-Bromage H, Tempst P, Sabatini DM. mTOR interacts with raptor to form a nutrient-sensitive complex that signals to the cell growth machinery. Cell 2002; 110:163–75.

20. Heuser JE, Reese TS. Evidence for recycling of synaptic vesicle membrane during transmitter release at the frog neuromuscular junction. J Cell Biol 1973; 57:315–44.

21. Saheki Y, De Camilli P. Synaptic vesicle endocytosis. Cold Spring Harb Perspect Biol 2012; 4:a005645.

22. Südhof TC. The synaptic vesicle cycle: a cascade of protein–protein interactions. Nature 1995; 375:645–53.

23. Condliffe SB, Corradini I, Pozzi D, Verderio C, Matteoli M. Endogenous SNAP-25 Regulates Native Voltage-gated Calcium Channels in Glutamatergic Neurons. J Biol Chem 2010; 285:24968–76.

24. Chen Y, Deng L, Maeno-Hikichi Y, Lai M, Chang S, Chen G, Zhang JF. Formation of an endophilin-Ca2+ channel complex is critical for clathrin-mediated synaptic vesicle endocytosis. Cell 2003; 115:37–48.

25. Sørensen JB, Matti U, Wei S-H, Nehring RB, Voets T, Ashery U, Binz T, Neher E, Rettig J. The SNARE protein SNAP-25 is linked to fast calcium triggering of exocytosis. Proc Natl Acad Sci U S A [Internet] 2002; 99:1627–32. Available from: http://www.ncbi.nlm.nih.gov/pmc/articles/PMC122241/

26. Maday S, Wallace KE, Holzbaur ELF. Autophagosomes initiate distally and mature during transport toward the cell soma in primary neurons. J Cell Biol 2012; 196:407–17.

27. Yang S, Park D, Manning L, Hill SE, Cao M, Xuan Z, Gonzalez I, Dong Y, Clark B, Shao L, et al. Presynaptic autophagy is coupled to the synaptic vesicle cycle via ATG-9. Neuron 2022; 110:824–840.e10.

28. Taoufiq Z, Ninov M, Villar-Briones A, Wang HY, Sasaki T, Roy MC, Beauchain F, Mori Y, Yoshida T, Takamori S, et al. Hidden proteome of synaptic vesicles in the mammalian brain. Proc Natl Acad Sci U S A 2020; 117:33586–96.

29. Murdoch JD, Rostosky CM, Gowrisankaran S, Arora AS, Soukup S-F, Vidal R, Capece V, Freytag S, Fischer A, Verstreken P, et al. Endophilin-A Deficiency Induces the Foxo3a-Fbxo32 Network in the Brain and Causes Dysregulation of Autophagy and the Ubiquitin-Proteasome System. Cell Rep 2016; 17:1071–86.

30. Okerlund ND, Schneider K, Leal-Ortiz S, Montenegro-Venegas C, Kim SA, Garner LC, Gundelfinger ED, Reimer RJ, Garner CC. Bassoon Controls Presynaptic Autophagy through Atg5. Neuron 2017; 93:897–913.e7.

31. Kononenko NL, Claßen GA, Kuijpers M, Puchkov D, Maritzen T, Tempes A, Malik AR, Skalecka A, Bera S, Jaworski J, et al. Retrograde transport of TrkB-containing autophagosomes via the adaptor AP-2 mediates neuronal complexity and prevents neurodegeneration. Nat Commun 2017; 8:14819.

32. McInnes J, Wierda K, Snellinx A, Bounti L, Wang Y-C, Stancu I-C, Apóstolo N, Gevaert K, Dewachter I, Spires-Jones TL, et al. Synaptogyrin-3 Mediates Presynaptic Dysfunction Induced by Tau. Neuron 2018; 97:823–835.e8.

33. Zhou L, McInnes J, Wierda K, Holt M, Herrmann AG, Jackson RJ, Wang Y-C, Swerts J, Beyens J, Miskiewicz K, et al. Tau association with synaptic vesicles causes presynaptic dysfunction. Nat Commun 2017; 8:15295.

34. Soukup S, Verstreken P. PIWIL1 protein power targets tau therapy. Nat Neurosci 2014; 17:334–5.

35. Frost B, Hemberg M, Lewis J, Feany MB. Tau promotes neurodegeneration through global chromatin relaxation. Nat Neurosci 2014; 17:357–66.

36. Nixon RA, Yang D-S. Autophagy failure in Alzheimer’s disease--locating the primary defect. Neurobiol Dis 2011; 43:38–45.

37. Lim F, Hernández F, Lucas JJ, Gómez-Ramos P, Morán MA, Avila J. FTDP-17 mutations in tau transgenic mice provoke lysosomal abnormalities and Tau filaments in forebrain. Mol Cell Neurosci 2001; 18:702–14.

38. Largo-Barrientos P, Apóstolo N, Creemers E, Callaerts-Vegh Z, Swerts J, Davies C, McInnes J, Wierda K, De Strooper B, Spires-Jones T, et al. Lowering Synaptogyrin-3 expression rescues Tau-induced memory defects and synaptic loss in the presence of microglial activation. Neuron 2021; 109:767–777.e5.

39. Hernandez-Diaz S, Ghimire S, Sanchez-Mirasierra I, Montecinos-Oliva C, Swerts J, Kuenen S, Verstreken P, Soukup S-F. Endophilin-B regulates autophagy during synapse development and neurodegeneration. Neurobiol Dis 2022; 163:105595.

40. Shehata M, Matsumura H, Okubo-Suzuki R, Ohkawa N, Inokuchi K. Neuronal stimulation induces autophagy in hippocampal neurons that is involved in AMPA receptor degradation after chemical long-term depression. J Neurosci 2012; 32:10413–22.

41. Wang T, Martin S, Papadopulos A, Harper CB, Mavlyutov TA, Niranjan D, Glass NR, Cooper-White JJ, Sibarita JB, Choquet D, et al. Control of autophagosome axonal retrograde flux by presynaptic activity unveiled using botulinum neurotoxin type A. J Neurosci 2015; 35:6179–94.

42. Stevens RJ, Akbergenova Y, Jorquera R a, Littleton JT. Abnormal synaptic vesicle biogenesis in Drosophila synaptogyrin mutants. J Neurosci 2012; 32:18054–67, 18067a.

43. Sanchez-Mirasierra I, Hernandez-Diaz S, Ghimire S, Montecinos-Oliva C, Soukup S-F. Macros to Quantify Exosome Release and Autophagy at the Neuromuscular Junction of Drosophila Melanogaster. Front Cell Dev Biol 2021; 9:1–13.

44. Hamada FN, Rosenzweig M, Kang K, Pulver SR, Ghezzi A, Jegla TJ, Garrity PA. An internal thermal sensor controlling temperature preference in Drosophila. Nature 2008; 454:217–20.

45. Pulver SR, Pashkovski SL, Hornstein NJ, Garrity P a, Griffith LC. Temporal dynamics of neuronal activation by Channelrhodopsin-2 and TRPA1 determine behavioral output in Drosophila larvae. J Neurophysiol 2009; 101:3075–88.

46. Yoshii M, Watabe S, Murashima YL, Nukada T, Shiotani T. Cellular mechanism of action of cognitive enhancers: Effects of nefiracetam on neuronal Ca2+ channels. In: Alzheimer Disease and Associated Disorders. 2000.

47. Yoshii M, Watabe S. Enhancement of neuronal calcium channel currents by the nootropic agent, nefiracetam (DM-9384), in NG108-15 cells. Brain Res 1994; 642:123–31.

48. Nishizaki T, Matsuoka T, Nomura T, Sumikawa K, Shiotani T, Watabe S, Yoshii M. Nefiracetam modulates acetylcholine receptor currents via two different signal transduction pathways. Mol Pharmacol 1998; 53:1–5.

49. Young JE, Martinez RA, La Spada AR. Nutrient deprivation induces neuronal autophagy and implicates reduced insulin signaling in neuroprotective autophagy activation. J Biol Chem 2009; 284:2363–73.

50. Bjedov I, Toivonen JM, Kerr F, Slack C, Jacobson J, Foley A, Partridge L. Mechanisms of life span extension by rapamycin in the fruit fly Drosophila melanogaster. Cell Metab 2010; 11:35–46.

51. Truckenbrodt S, Viplav A, Jähne S, Vogts A, Denker A, Wildhagen H, Fornasiero EF, Rizzoli SO. Newly produced synaptic vesicle proteins are preferentially used in synaptic transmission. EMBO J 2018; 37:e98044.

52. Juan CA, Pérez de la Lastra JM, Plou FJ, Pérez-Lebeña E. The Chemistry of Reactive Oxygen Species (ROS) Revisited: Outlining Their Role in Biological Macromolecules (DNA, Lipids and Proteins) and Induced Pathologies. Int J Mol Sci 2021; 22.

53. Hafner AS, Donlin-Asp PG, Leitch B, Herzog E, Schuman EM. Local protein synthesis is a ubiquitous feature of neuronal pre- And postsynaptic compartments. Science (80-) 2019; 364.

54. Nagy P, Kárpáti M, Varga A, Pircs K, Venkei Z, Takáts S, Varga K, Erdi B, Hegedűs K, Juhász G. Atg17/FIP200 localizes to perilysosomal Ref(2)P aggregates and promotes autophagy by activation of Atg1 in Drosophila. Autophagy 2014; 10:453–67.

55. Matoba K, Kotani T, Tsutsumi A, Tsuji T, Mori T, Noshiro D, Sugita Y, Nomura N, Iwata S, Ohsumi Y, et al. Atg9 is a lipid scramblase that mediates autophagosomal membrane expansion. Nat Struct Mol Biol 2020; 27:1185–93.

56. Papinski D, Schuschnig M, Reiter W, Wilhelm L, Barnes C a, Maiolica A, Hansmann I, Pfaffenwimmer T, Kijanska M, Stoffel I, et al. Early steps in autophagy depend on direct phosphorylation of Atg9 by the Atg1 kinase. Mol Cell 2014; 53:471–83.

57. Yamamoto H, Kakuta S, Watanabe TM, Kitamura A, Sekito T, Kondo-Kakuta C, Ichikawa R, Kinjo M, Ohsumi Y. Atg9 vesicles are an important membrane source during early steps of autophagosome formation. J Cell Biol 2012; 198:219–33.

58. Binotti B, Ninov M, Cepeda AP, Ganzella M, Matti U, Riedel D, Urlaub H, Sambandan S, Jahn R. ATG9 resides on a unique population of small vesicles in presynaptic nerve terminals. Autophagy 2024; 20:883–901.

59. Kallergi E, Siva Sankar D, Matera A, Kolaxi A, Paolicelli RC, Dengjel J, Nikoletopoulou V. Profiling of purified autophagic vesicle degradome in the maturing and aging brain. Neuron 2023; 111:2329–2347.e7.

60. Zhang B, Higuchi M, Yoshiyama Y, Ishihara T, Forman MS, Martinez D, Joyce S, Trojanowski JQ, Lee VM-Y. Retarded axonal transport of R406W mutant tau in transgenic mice with a neurodegenerative tauopathy. J Neurosci Off J Soc Neurosci 2004; 24:4657–67.

61. Feng Q, Luo Y, Zhang X-N, Yang X-F, Hong X-Y, Sun D-S, Li X-C, Hu Y, Li X-G, Zhang J-F, et al. MAPT/Tau accumulation represses autophagy flux by disrupting IST1-regulated ESCRT-III complex formation: a vicious cycle in Alzheimer neurodegeneration. Autophagy 2020; 16:641–58.

62. Wang Y, Martinez-Vicente M, Krüger U, Kaushik S, Wong E, Mandelkow EM, Cuervo AM, Mandelkow E. Tau fragmentation, aggregation and clearance: The dual role of lysosomal processing. Hum Mol Genet 2009; 18:4153–70.

63. Egaña LA, Cuevas RA, Baust TB, Parra LA, Leak RK, Hochendoner S, Peña K, Quiroz M, Hong WC, Dorostkar MM, et al. Physical and functional interaction between the dopamine transporter and the synaptic vesicle protein synaptogyrin-3. J Neurosci Off J Soc Neurosci 2009; 29:4592–604.

64. Hernandez D, Torres C a, Setlik W, Cebrián C, Mosharov E V, Tang G, Cheng H-C, Kholodilov N, Yarygina O, Burke RE, et al. Regulation of presynaptic neurotransmission by macroautophagy. Neuron 2012; 74:277–84.

65. Juhász G, Hill JH, Yan Y, Sass M, Baehrecke EH, Backer JM, Neufeld TP. The class III PI(3)K Vps34 promotes autophagy and endocytosis but not TOR signaling in Drosophila. J Cell Biol 2008; 181:655–66.

66. Wittmann CW, Wszolek MF, Shulman JM, Salvaterra PM, Lewis J, Hutton M, Feany MB. Tauopathy in Drosophila: neurodegeneration without neurofibrillary tangles. Science 2001; 293:711–4.

67. Kiss V, Jipa A, Varga K, Takáts S, Maruzs T, Lőrincz P, Simon-Vecsei Z, Szikora S, Földi I, Bajusz C, et al. Drosophila Atg9 regulates the actin cytoskeleton via interactions with profilin and Ena. Cell Death Differ 2020; 27:1677–92.

68. Guruharsha KG, Rual J-F, Zhai B, Mintseris J, Vaidya P, Vaidya N, Beekman C, Wong C, Rhee DY, Cenaj O, et al. A protein complex network of Drosophila melanogaster. Cell 2011; 147:690–703.

69. Nagy P, Kárpáti M, Varga A, Pircs K, Venkei Z, Takáts S, Varga K, Erdi B, Hegedűs K, Juhász G. Atg17/FIP200 localizes to perilysosomal Ref(2)P aggregates and promotes autophagy by activation of Atg1 in Drosophila. Autophagy 2014; 10:453–67.

70. Barroso-Gomila O, Trulsson F, Muratore V, Canosa I, Merino-Cacho L, Cortazar AR, Pérez C, Azkargorta M, Iloro I, Carracedo A, et al. Identification of proximal SUMO-dependent interactors using SUMO-ID. Nat Commun 2021; 12:6671.

