## supplemental information for "Synaptogyrin regulates neuronal activity-induced autophagy to degrade synaptic vesicle components and pathological Tau"

**Supplement Material and Methods**

**Detailed genotype description.**

The following genotypes were used in this study as followed:

*w;; UAS- mCherry::Atg8/D42-Gal4*

*w; syngr$^{1}$; UAS- mCherry::Atg8/D42-Gal4*

*w; syngr $^{1}$; UAS- syngr::myc /D42-Gal4, UAS- mCherry::Atg8*

*w; syngr $^{1}$; UAS-mCherry-GFP::Atg8/D42-Gal4*

*w; UAS-TrpA1/ UAS-Atg8mCherry, OK6-Gal4*

*w; UAS-TrpA1/ UAS-mCherry::Atg8, OK6-Gal4;UAS- syngr::myc/+*

*w; UAS- mCherry::Atg8, OK6-Gal4/+; UAS- syngr$^{shRNA}$ /+*

*w; UAS- mCherry::Atg8, OK6-Gal4/+; UAS-endoB::HA/UAS- syngr$^{shRNA}$*

*w ;UAS-Atg8mCherry, OK6-Gal4/+*

*w;; UAS-mCherry-GFP::Atg8/D42-Gal4*

*w; UAS- mCherry::Atg8, OK6-Gal4/+*

*w; UAS- mCherry::Atg8, OK6-Gal4/+; UAS-endoB::HA/+*

*w; UAS- mCherry::Atg8, OK6-Gal4/+; UAS- syngr::myc/+*

*w; UAS- mCherry::Atg8, OK6-Gal4/+; UAS- syngr::myc/UAS-endoB::HA*

*w; endoB$^{4}$,UAS- mCherry::Atg8/endoB$^{4}$;D42-GAL4/ UAS-TrpA1*

*w; UAS- mCherry::Atg8/+;D42-GAL4/ UAS-TrpA1*

*w;; UAS- syngr$^{shRNA}$ / UAS- mCherry::Atg8, D42-GAL4*

*w; endoB$^{4}$; UAS- syngr$^{shRNA}$ / UAS- mCherry::Atg8, D42-GAL4*

*w*

*w; syngr$^{1}$*

*w; syngr$^{1}$; atg17$^{d130}$*

*w;; atg17$^{d130}$*

*w; atg9$^{9B5}$; HA::atg9*

*w; atg9$^{9B5}$*

*w;;UAS-Mito-GFP, DA-Gal4/+*

*w; GFP::Atg9*

*w; UAS- mCherry::Atg8/+;UAS- GFP::Atg9, D42-GAL4/+*

*shi$^{ts1}$;; UAS- GFP::Atg9, D42-GAL4/+*

*w;; UAS- GFP::Atg9, D42-GAL4/+*

*w; UAS- mCherry::Atg8, OK6-Gal4/+; UAS-hTau$^{R406W}$/+*

*w;; UAS- mCherry-GFP::Atg8/ UAS-hTau$^{R406W}$, D42-Gal4*

*w; gyrin$^{1}$/+; UAS- mCherry-GFP::Atg8/ UAS-hTau$^{R406W}$, D42-Gal4*

*w;; UAS-hTau$^{R406W}$, D42-Gal4/+*

*w; gyrin$^{1}$/+; UAS-hTau$^{R406W}$, D42-Gal4/+*

*w; gyrin$^{1}$/+; UAS-hTau$^{R406W}$, D42-Gal4/atg17$^{d130}$*

*w;; UAS- mCherry::Atg8/ UAS-hTau$^{R406W}$, D42-Gal4*

*w; gyrin$^{1}$/+; UAS- mCherry::Atg8/ UAS-hTau$^{R406W}$, D42-Gal4*

*w; atg9$^{9B5}$; UAS- syngr$^{shRNA}$ / UAS- mCherry::Atg8, D42-GAL4*

*w; atg9$^{9B5}$; UAS- mCherry::Atg8, D42-GAL4/+*

*w; gyrin$^{1}$ ; UAS- GFP::Atg9, D42-GAL4/+*

*w;;UAS- GFP::Atg9, D42-GAL4/+*

*w;;UAS- GFP::Atg9, D42-GAL4/ UAS- syngr::myc*

*w*; *endoB*$^{4}$,UAS- mCherry::Atg8/*endoB*$^{4}$;D42-Gal4/+

*w*; *endoB*$^{4}$,UAS-mCherry::Atg8/*endoB*$^{4}$;D42-GAL4/*endoB*$^{Bac}$

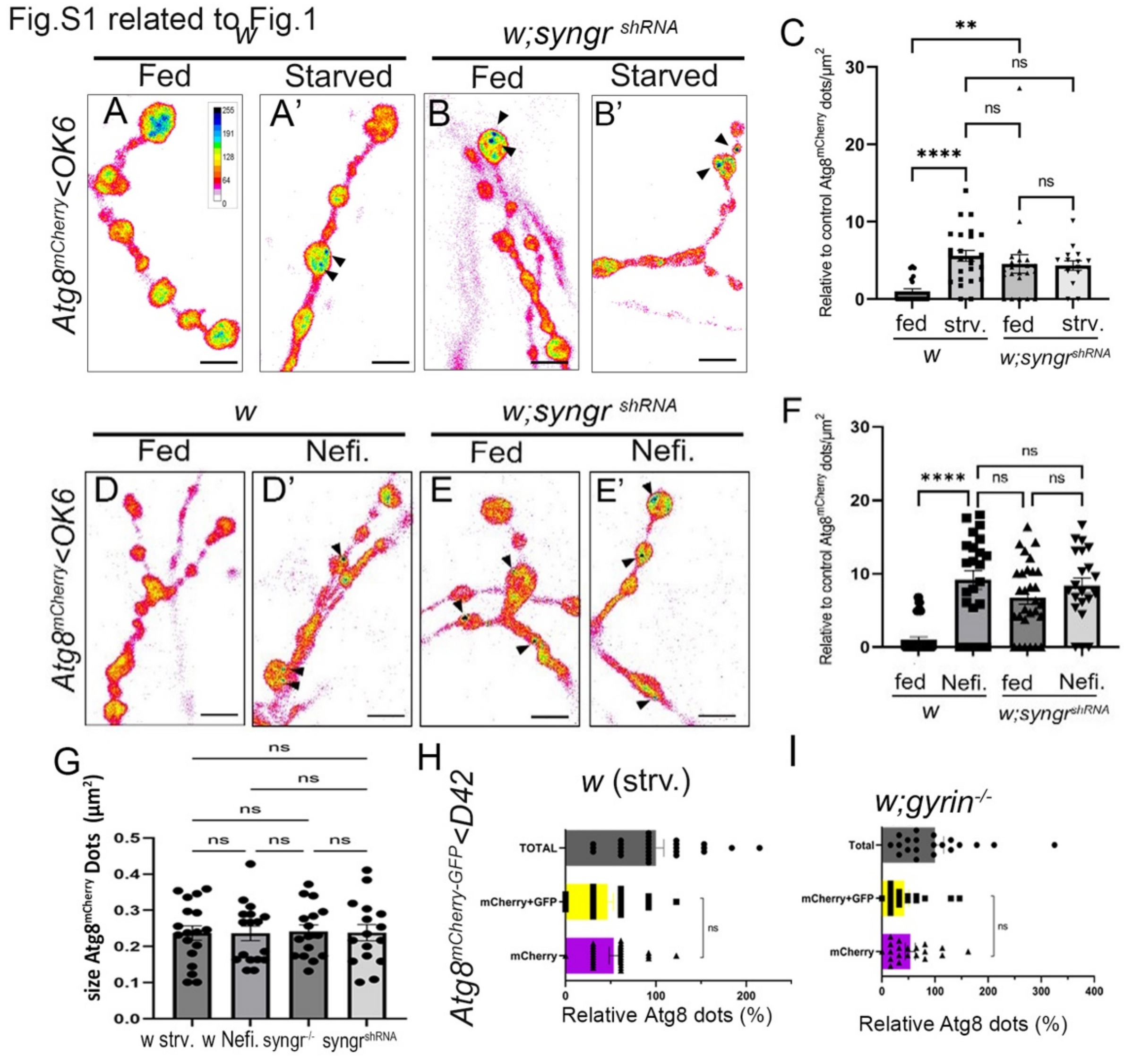


**Figure S1. Neuron specific knockdown of *synaptogyrin* leads to increased autophagy levels at the Drosophila NMJ synapse, Related to Fig. 1. (A-B')** Imaging of autophagy marker Atg8$^{mCherry}$ in fed **(A,B)** or 4h starved **(A',B')** wildtype animals or animals expressing shRNA against *syngr* in motor neurons (*syngr*$^{shRNA}$). Fluorescence intensities are shown using the indicated scale in **(A)**. **(C)** Quantification of number of Atg8$^{mCherry}$ dots at the NMJ shows significant increased amount of autophagosomes in animals with neuronal knock down of *syngr*. n >24 individual synapses from a minimum of 5 larvae.

**(D-E')** Imaging of autophagy marker $Atg8^{mCherry}$ in **(D,E)** non-stimulated animals incubated for 30 mins in HL3 containing $CaCl_2$ and **(D', E')** animals incubated for 30 mins in HL3 containing calcium channel agonist Nefiracetam (Nefi) and $CaCl_2$ to induce synaptic autophagy. **(F)** Quantification of the number of $Atg8^{mCherry}$ dots at presynaptic terminals. n >24 individual synapses from a minimum of 5 larvae. Scale bars = 5 µm.

**(G)** Quantification of $Atg8^{mCherry}$ dots size at presynaptic terminals in 4h starved wildtype animals, wildtype animals treated with Nefi, *syngr* null mutant and animals with neuronal knock down of *syngr* shows no statistically significant difference.

**(H)** Quantification of live imaging of autophagy marker Atg8mCherry-GFP to assess autophagic flux in 4 h starved wildtype animals, shows total number of Atg8 (GFP+mCherry plus mCherry alone) dots and the relative portion of white (GFP+ mCherry) and magenta (mCherry) Atg8 dots in percent (representative images are shown in Figure 1F-F"). **(I)** Quantification of live imaging of *syngr* null mutants expressing Atg8mCherry-GFP to analyze autophagic flux shows total number of Atg8 (GFP+mCherry plus mCherry alone) dots and the relative portion of white (GFP+mCherry) and magenta (mCherry) Atg8 dots in percent (representative images are shown in Figure 1G-G").

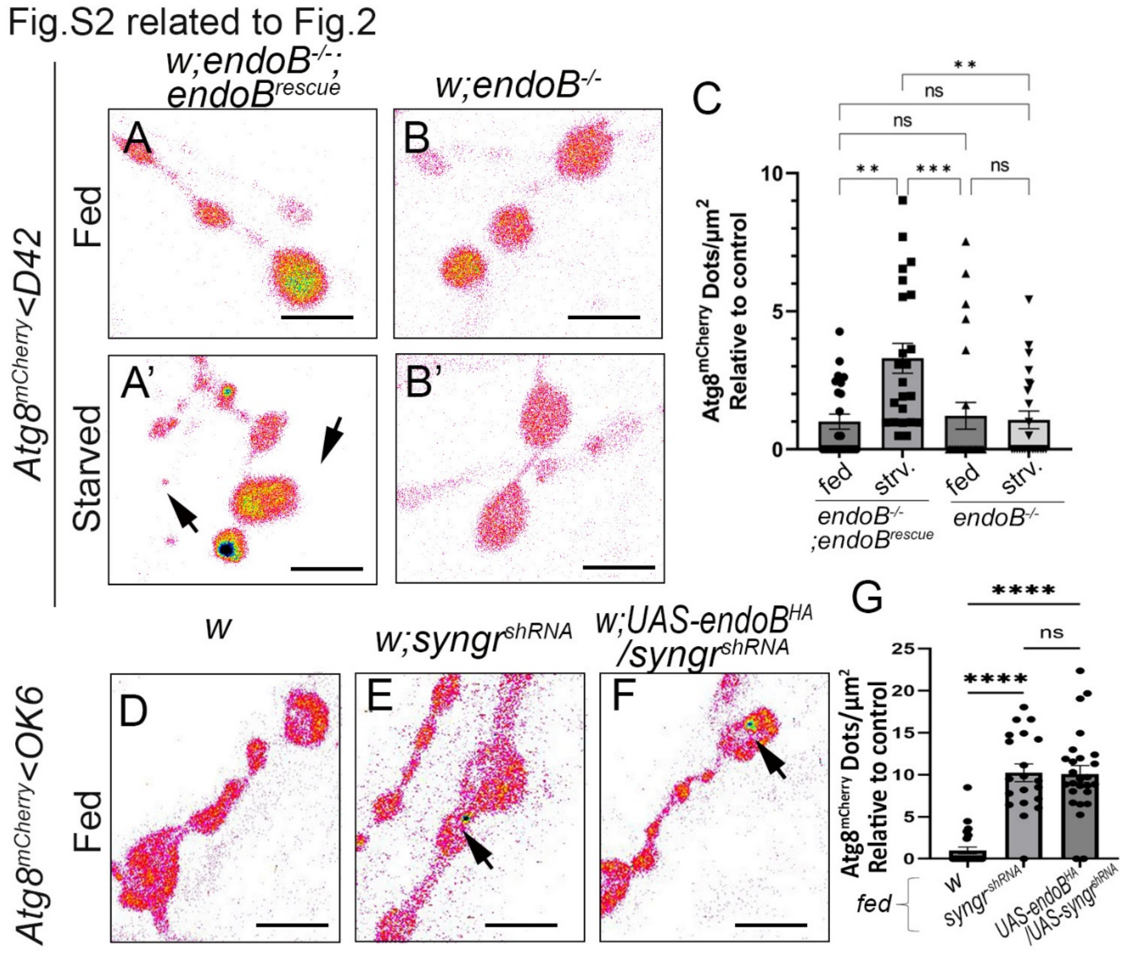


**Figure S2, Related to Fig. 2. EndoB function is required for starvation induced autophagy at the presynaptic terminal. (A-B')** Neuromuscular junction (NMJ) boutons expressing Atg8$^{mCherry}$ using a motor neuron driver (D42-Gal4). **(A,A')** *endoB* genomic recues construct in *endoB$^{4}$* mutants reestablishes number of autophagosomes after starvation (arrows). **(B,B')** *endoB$^{4}$* null mutants (*endoB$^{-/-}$*) do not show increase in number of autophagosome after starvation. **(C)** Statistical significant increased amount of autophagosomes in *endoB$^{4}$* mutants with *endoB* genomic recues construct compared to *endoB*$^{4}$ mutants alone. n >20 individual synapses from a minimum of 5 larvae.

Imaging of wildtype fed **(D)**, fed animals expressing shRNA against *syngr* in motor neurons **(E)** and in animals expressing shRNA against *syngr* and overexpressing *endoB*$^{HA}$ **(F)**. **(G)** Quantification of Atg8 positive dots show no significant differences between animals with neuronal knockdown of *syngr* and aimals with neuronal knockdown of *syngr* and overexpression of *endoB*$^{HA}$. Scale bars= 5 μm

Fig.S3 related to Fig.3

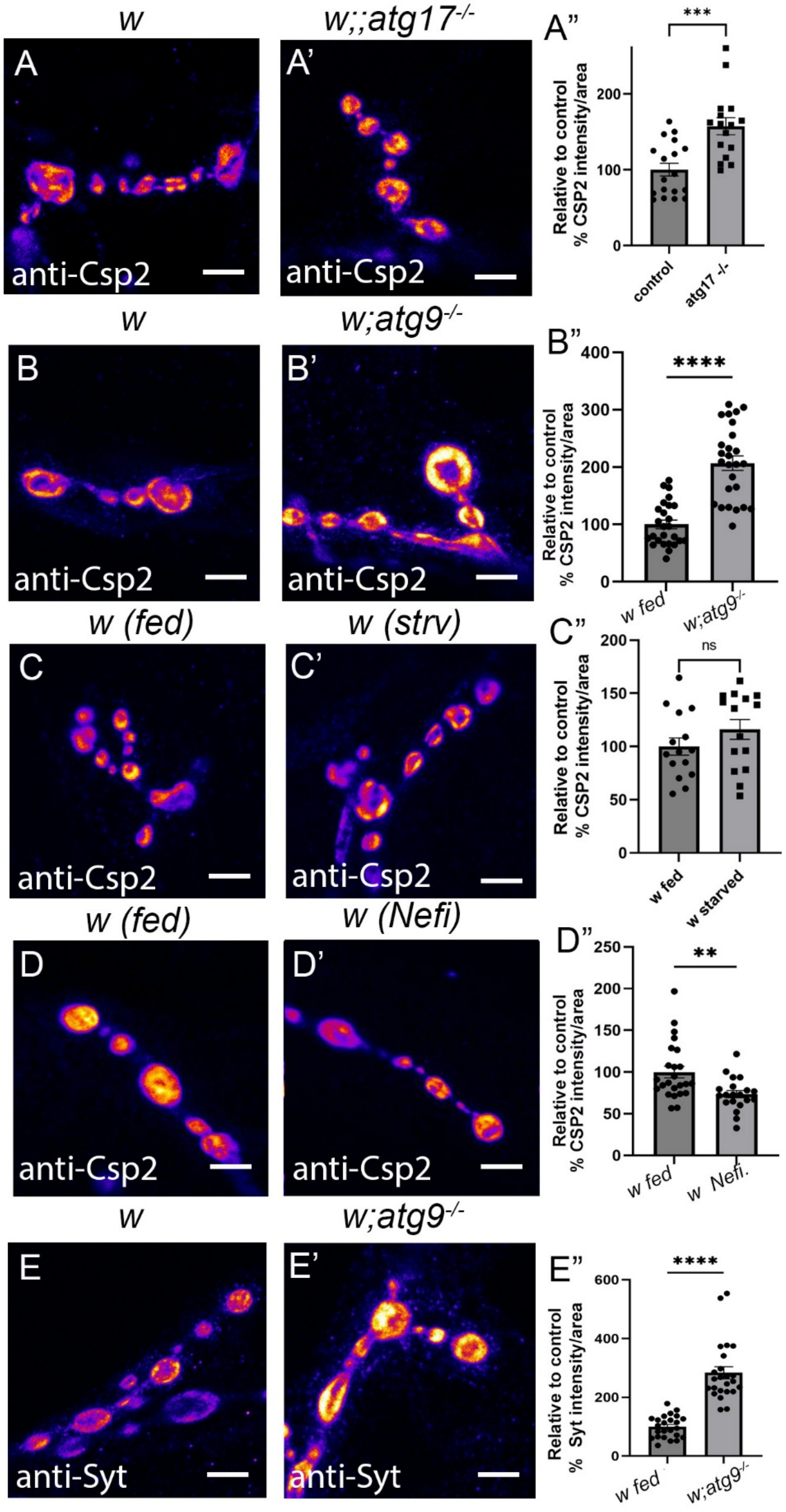

**Figure S3., Related to Fig. 3. Degradation of synaptic vesicle protein Csp2 by autophagy. (A,A')** Immunostainings of synaptic vesicle component Csp2 in **(A)** wildtype and **(A')** *atg17* null mutant (w;;atg17$^{-/-}$) NMJs. **(A'')** Quantification of fluorescence intensities indicates a significant increase of the synaptic vesicle marker Csp2 in *atg17* null mutants compared to wildtype.

Immunostainings of Csp2 in **(B)** fed wildtype and **(B')** *atg9* null mutant (w;;atg9$^{-/-}$) NMJs. **(B'')** Quantification of fluorescence intensities indicates a significant increase of the Csp2 in *atg9* null mutants compared to wildtype.

**(C,C')** Csp2 immunostaining in synapses of fed and 4h starved larvae. **(C'')** Quantification of fluorescence intensities indicates no significant difference of the synaptic vesicle marker Csp2 in fed wildtype animals compared to starved wildtype animals.

Immunostainings of Csp2 in **(D)** fed wildtype NMJs and **(D')** wildtype animals incubated for 30 min with Nefiracetam (Nefi) to induce neuronal activity-induced autophagy in NMJs. **(D'')** Quantification of fluorescence intensities indicates a significant decrease of Csp2 after induction of neuronal activity-induced autophagy compared to wildtype.

Immunostainings of synaptic vesicle marker Synaptotagmin (Syt) in **(E)** fed wildtype and **(E')** *atg9* null mutant (w;;atg9$^{-/-}$) NMJs. **(E'')** Quantification of fluorescence intensities indicates a significant increase of the Syt in *atg9* null mutants compared to wildtype.

**(A-E'')** n >22 individual synapses from a minimum of 5 larvae. Scale bars= 5µm

Fig.S4 related to Fig.4

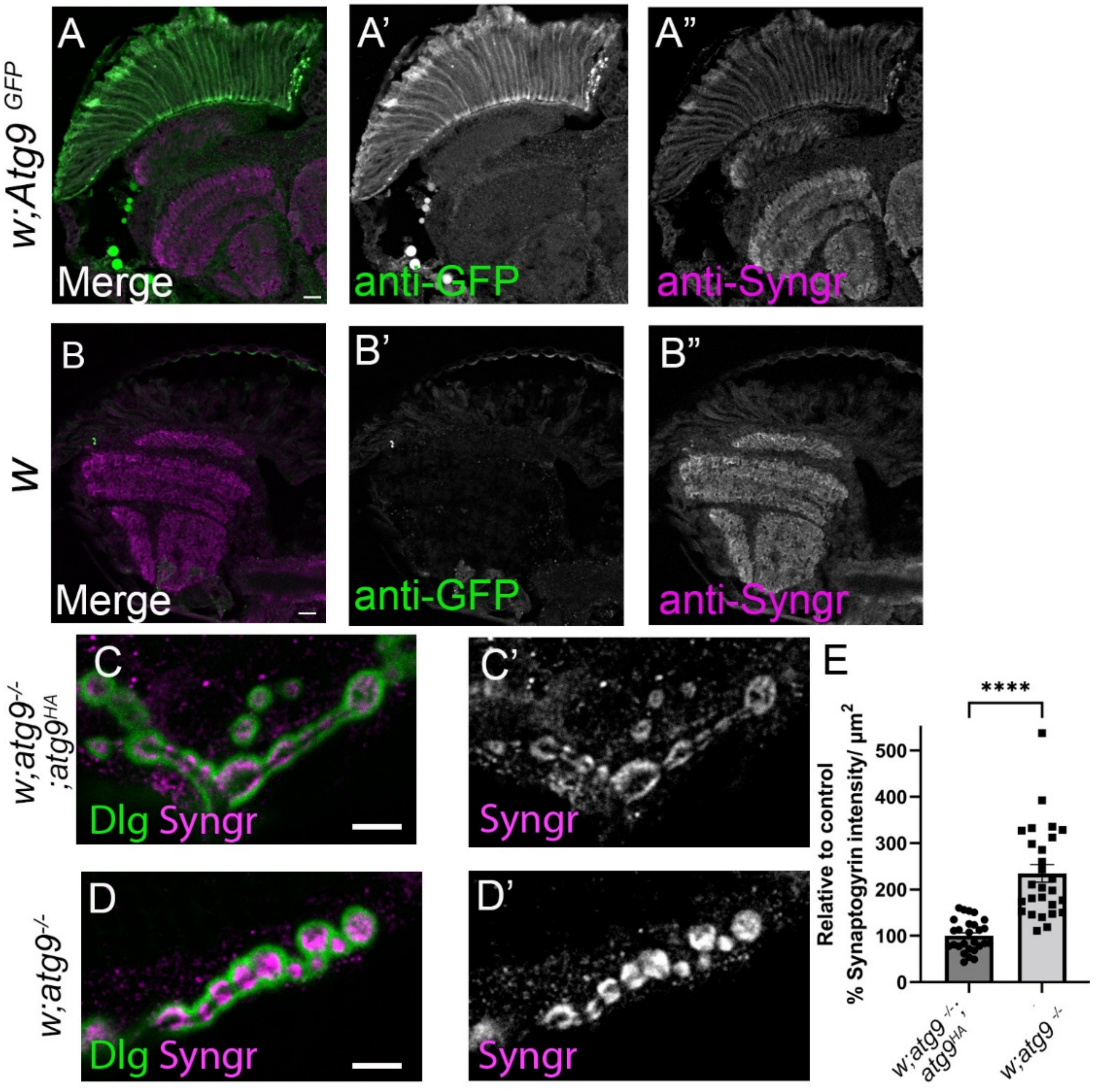


**Figure S4. Atg9 and Synaptogyrin localize to synaptic structures in the adult visual system, Related to Fig. 4. (A-A")** Longitudinal section of 5-day-old adult flies with endogenous expression of Atg9::GFP (green) in the visual system **(A').** **(B-B")** Longitudinal section of 5-day-old wildtype adult flies shows only residual GFP signal **(B')**. Syngr expression (magenta) is similar in the visual system of flies expressing endogenous Atg9::GFP and wildtype flies **(A",B"). (C-D')** Labelling of

NMJ boutons of flies expressing *atg9*$^{HA}$ under endogenous levels in *atg9* mutant background **(C,C')** and of *atg9* null mutants **(D,D')** with anti-DLG (green) to mark the outline of the NMJ synapse and anti-Syngr (magenta). **(E)** Quantification shows a statistically significant increase of Syngr labelling in *atg9* null mutant compared to *atg9* null mutants with genomic rescue construct. Scale bars = 5μm. n >16 individual synapses from a minimum of 5 larvae.

Fig.S5

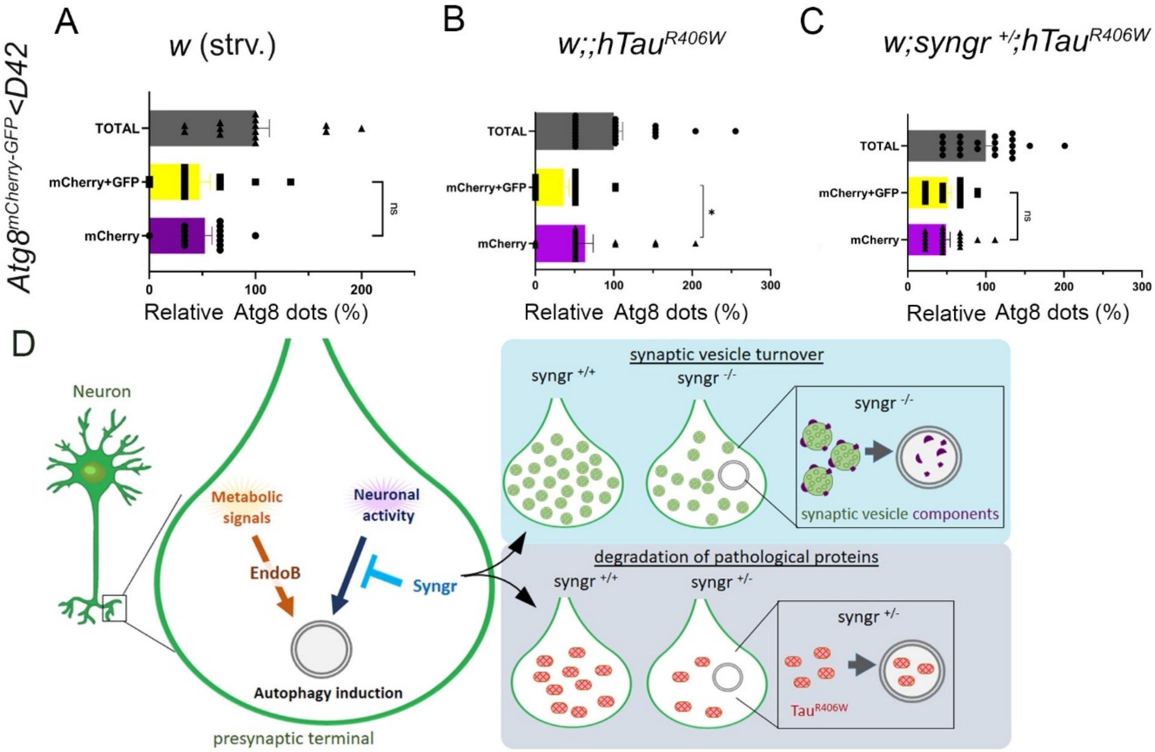


**Figure S5. Live imaging of Atg8$^{mCherry-GFP}$ to analyze autophagic flux in NMJ boutons. Related to Fig. 5.** Quantification of live imaging of autophagy marker Atg8mCherry-GFP to assess autophagic flux in starved wildtype flies **(A)**, flies expressing human hTau$^{R406W}$ **(B)** and of flies expressing hTau$^{R406W}$ with reduced levels of Syngr (*syngr* $^{-/+}$) **(C)**, shows total number of Atg8 (GFP+mCherry plus mCherry alone) dots and the relative portion of white (GFP+mCherry) and magenta (mCherry) Atg8 dots in percentage (n >27 individual synapses from a minimum of 5 larvae) in NMJ boutons of starved wildtype flies **(A)**, flies expressing hTau$^{R406W}$ **(B)** and of flies expressing hTau$^{R406W}$ with reduced levels of Syngr (*syngr* $^{-/+}$) **(C).** Representative images are shown in Figure 5 **D-F”**. **(D)** Cartoon summarizes Syngr function as negative regulator specifically for neuronal-activity induced autophagy to turnover synaptic vesicle components and phatological proteinsthat associate to synaptic vesicles like Tau.
